# PEPstrMOD2: Next-generation tertiary structure prediction of chemically modified and non-natural peptides

**DOI:** 10.64898/2026.06.22.733733

**Authors:** Saloni Jain, Naman Kumar Mehta, Sahil Raina, Pankaj Kumar, Varun, Gajendra P. S. Raghava

## Abstract

While most existing methods are limited to predicting the tertiary structures of proteins containing only canonical residues, the PEPstrMOD server (developed in 2015) pioneered structure prediction for chemically modified and non-natural peptides. Despite its widespread use, the original framework was restricted to peptides of 7 to 25 residues and relied on older backbone-prediction algorithms. To address these limitations, we present PEPstrMOD2, which introduces three major advancements over its predecessor. First, it replaces the original in-house coordinate generation with state-of-the-art deep learning (DL) algorithms, leveraging AlphaFold2 and ESMFold for highly accurate initial structure prediction. Secondly, it greatly expands the accessible chemical space through incorporation of new, AMBER force-field compatible library of 257 post-translational modifications (PTMs), 428 non-canonical amino acids (NCAAs), and 243 terminal modifications. Lastly, through the application of native scalability of AlphaFold2 (AF2) and ESMFold (EF), PEPstrMOD2 eliminates the original restrictions of the length, enabling the structural modeling of longer, complex therapeutic peptides and small proteins. We evaluated the performance of PEPstrMOD2 against state-of-the-art methods across three distinct peptide datasets. For the AfCyc dataset consisting of 80 cyclic peptides, PEPstrMOD2 obtained a competitive average atom-level Root Mean Square Deviation (RMSD) of 2.05 Å, compared to 1.13 Å by AlphaFold3 (AF3) and 1.82 Å by AfCycDesign. Remarkably, for the modified peptide ModPep433 dataset, PEPstrMOD2 outperformed AF3, achieving the lower average RMSD score of 4.49 Å against 4.67 Å of AF3. Furthermore, in the case of the ModPep16 benchmark, PEPstrMOD2 achieved 2.50 Å average RMSD value, which is two times more accurate than that of the original PEPstrMOD (5.84 Å). In summary, PEPstrMOD2 provides a powerful, high-throughput, and highly accurate platform to facilitate peptide-based drug development and structural biology research. While the original PEPstrMOD was restricted to a web server interface, PEPstrMOD2 is available as both an intuitive webserver and a standalone command-line tool via GitHub, featuring Docker support for easy deployment and reproducible, large-scale modeling pipelines (https://webs.iiitd.edu.in/raghava/pepstrmod/).

**Highlights:**

- PEPstrMOD2 predicts structures of chemically modified peptides using deep learning and molecular dynamics refinement.
- Supports 928 chemical modifications through custom AMBER based force-field libraries including PTMs, NCAAs, terminal, D- as well as cyclic modifications.
- Removes peptide length restrictions through AlphaFold2 and ESMFold integration.
- utperforms PEPstrMOD and is competitive with AF3 on different benchmark datasets.
- Available as both a web server and a standalone platform via GitHub and Docker.

**Author’s Biography:**

1. Saloni Jain is currently working as Ph.D. in Computational biology from Department of Computational Biology, Indraprastha Institute of Information Technology, New Delhi, India
2. Naman Kumar Mehta is currently working as Ph.D. in Computational biology from Department of Computational Biology, Indraprastha Institute of Information Technology, New Delhi, India
3. Sahil Raina is currently working as Ph.D. in Computational biology from Department of Computational Biology, Indraprastha Institute of Information Technology, New Delhi, India
4. Pankaj Kumar is currently working as Ph.D. in Computational biology from Department of Computational Biology, Indraprastha Institute of Information Technology, New Delhi, India
5. Varun is currently an integrated BS-MS student at Indian Institute of Science Education and Research (IISER) Pune, India. He is currently working as an intern on a project position at Department of Computational Biology, Indraprastha Institute of Information Technology (IIIT), New Delhi, India.
6. Gajendra P. S. Raghava is currently working as a Professor in the Department of Computational Biology, Indraprastha Institute of Information Technology, New Delhi, India.

## Introduction

Peptides have proven to be very important therapeutic compounds due to their target specificity, biological potency, and safer safety profile compared with traditional small molecule drugs [1]. They are used in diagnostics, drug delivery, and treatment of different diseases, and the number of peptide drugs that have been approved is growing together with the increasing number of peptide drugs under development [1–3]. Since the biological activity of peptides is based on their three-dimensional structure, knowing the structure is very important in understanding their mechanism of action and for designing drugs rationally. [4]. However, experimental structure determination by NMR spectroscopy or X-ray crystallography remains costly and time-intensive, restricting its use in large-scale peptide characterisation. This has driven sustained interest in computational methods capable of predicting peptide tertiary structures directly from sequence.

Over the past two decades, computational peptide structure prediction has progressed through several methodological generations (Table 1). The early methods used conformational sampling and energy minimization to predict the low-energy peptide structures. Geocore involved an extensive conformational search with an approximate energy function for structure prediction of small natural peptides, which could even accommodate disulfide-constrained peptides although it was computationally expensive for longer sequences [5]. The development of the β-turn prediction methods greatly improved the ability of the identification of the secondary structures in the peptides and opened the way for the subsequent peptide structure prediction algorithms <u>(Kaur and Raghava 2003; Kaur and Raghava 2004)</u>. Building upon this work, PEPstr used predicted beta-turns and secondary structures to generate initial conformations of the backbone, which were further refined through molecular dynamics (MD) simulations based on AMBER [6]. The Generalised Pattern Search algorithm similarly leveraged secondary structure predictions alongside an all-atom energy model and pattern search optimisation to identify near-native conformations of small bioactive peptides [7]. PEP-FOLD introduced a fundamentally different de novo framework using a Hidden Markov Model (HMM)-derived structural alphabet and coarse-grained force field, enabling rapid structure prediction of linear natural peptides and miniproteins directly from sequence without relying on secondary structure predictions [8]. Successive versions refined this approach: PEP-FOLD2 improved sampling efficiency and incorporated user-defined structural constraints, including disulfide bonds, while PEP-FOLD3 and PEP-FOLD4 extended sequence length coverage and introduced pH- and ionic strength-dependent modelling [9].

**Table 1.**
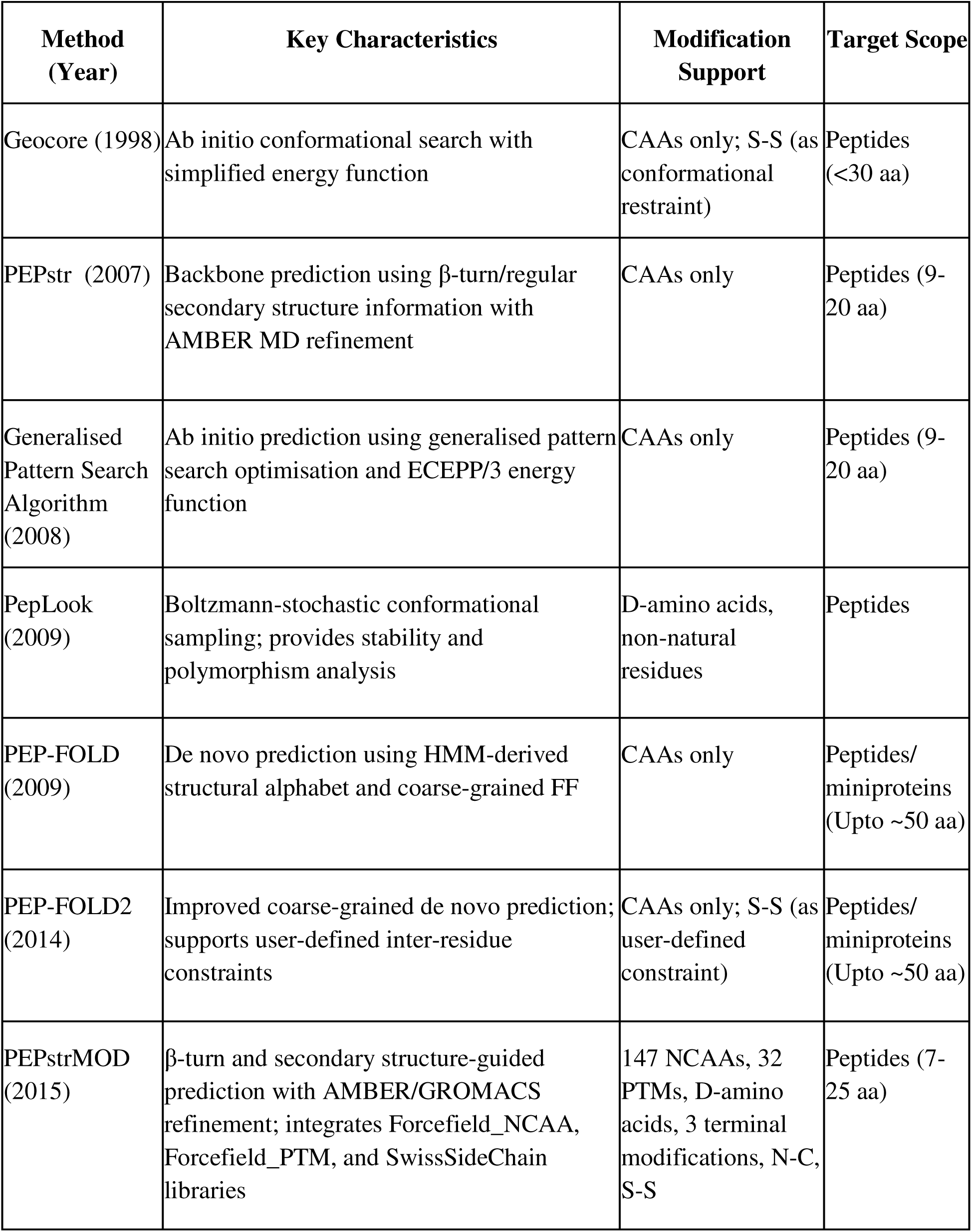
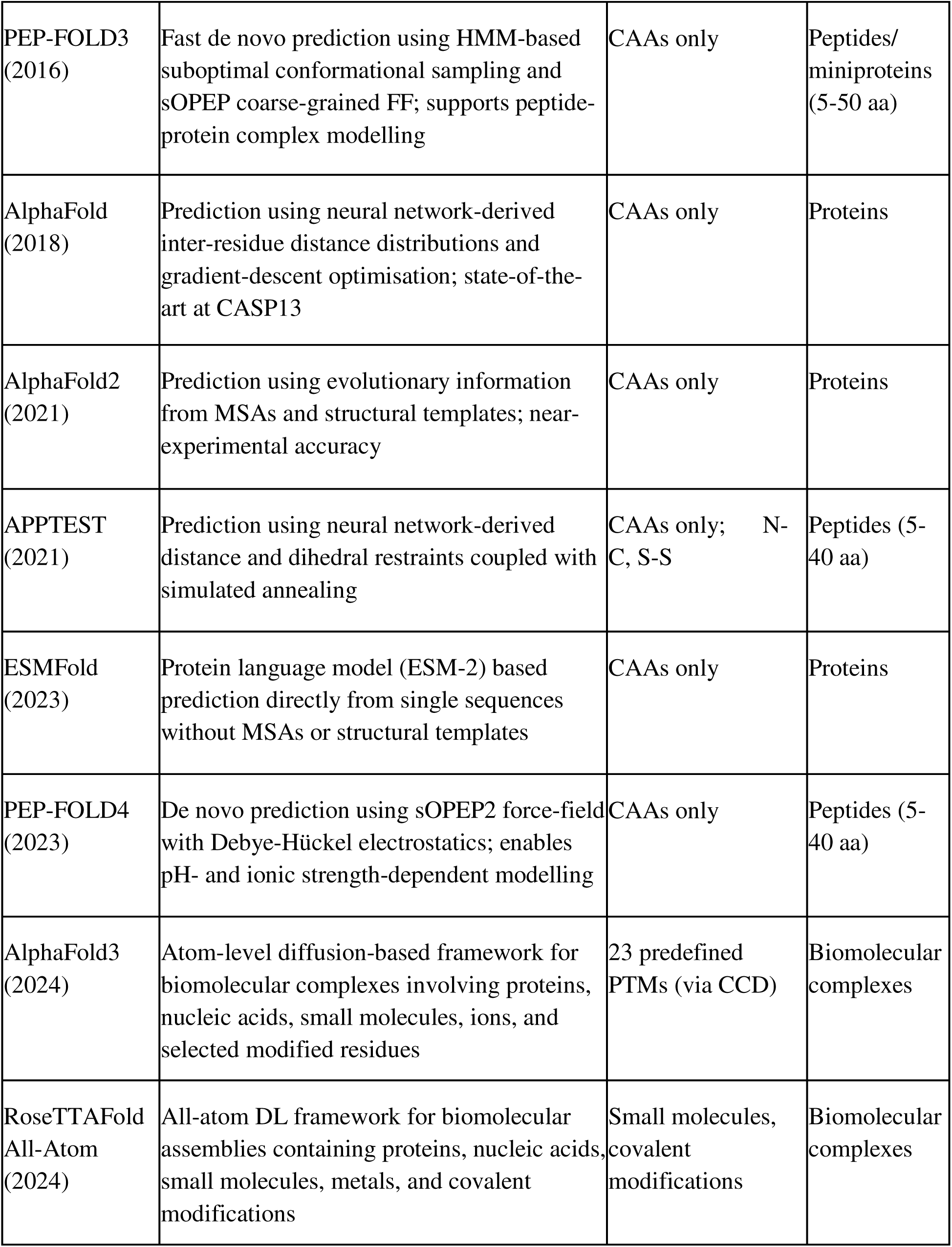
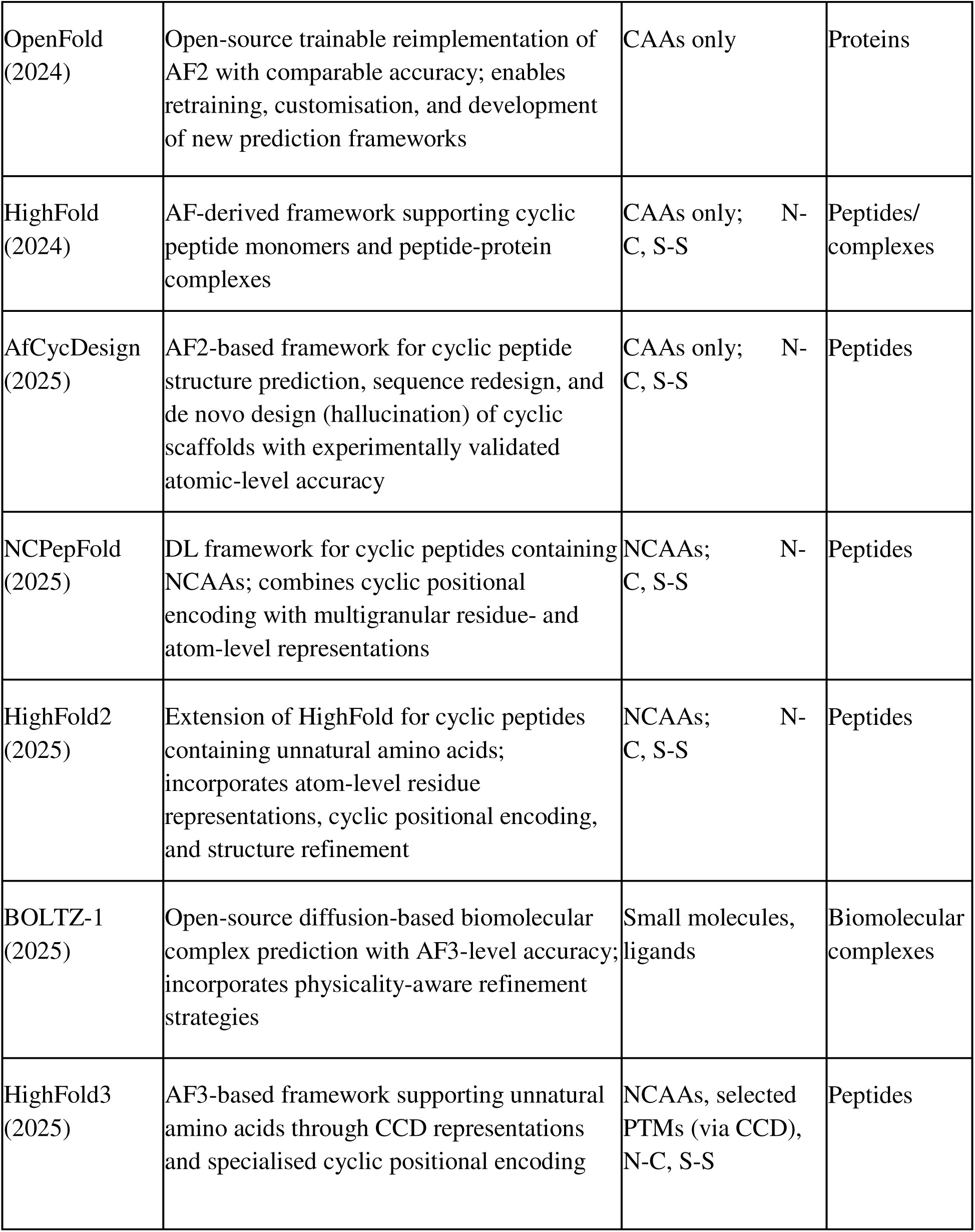
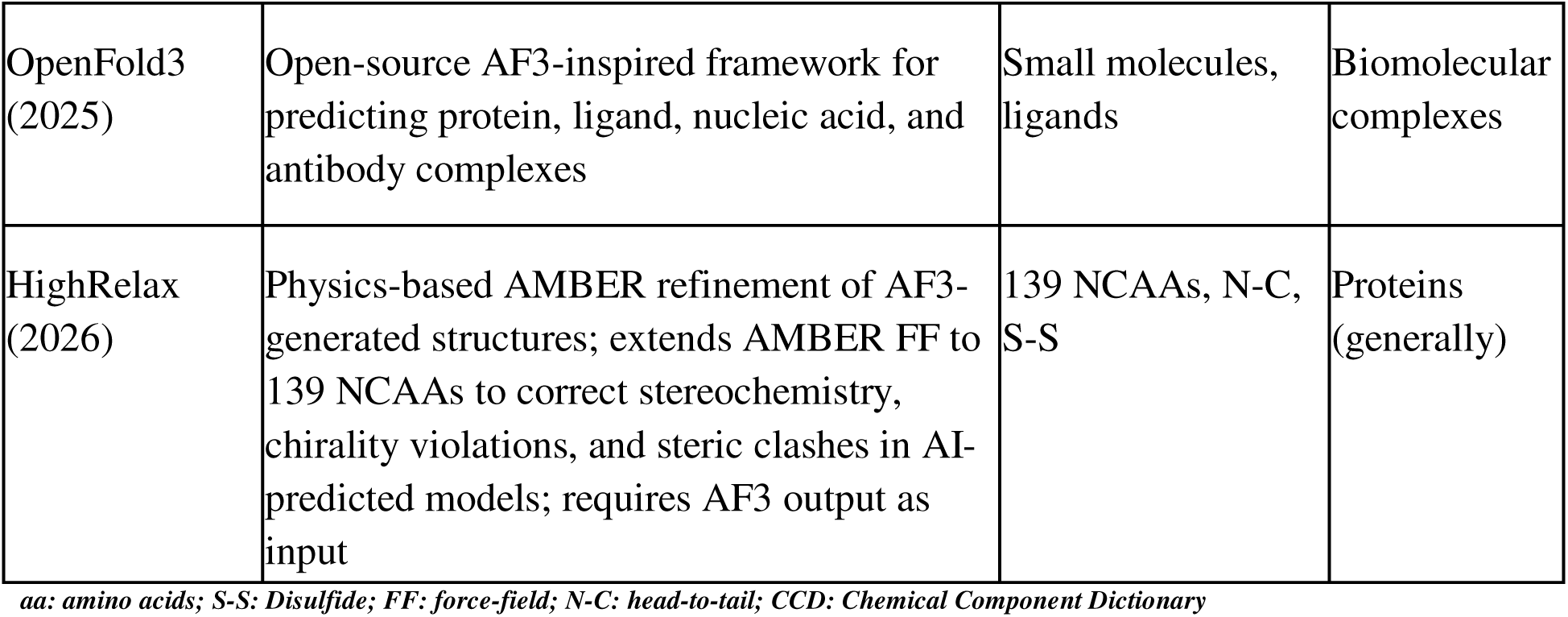
Evolution of computational methods for peptide/protein structure prediction and their key characteristics.

Although the mentioned techniques set the basis of computational peptide structure prediction, they were developed mainly for peptides that are constructed with canonical amino acids (CAAs). Nowadays, many peptides used in medicinal chemistry have different kinds of chemical modifications to enhance their pharmaceutical characteristics [10–17]. The other development line dealt with chemically modified peptides. PepLook was one of the first techniques that supported D-amino acids and non-canonical amino acid residues via Boltzmann stochastic sampling method for peptide polymorphism and stability modeling [18]. Further developments gave the chance to extend PEPstr algorithm by introducing Forcefield_NCAA [12], Forcefield_PTM [19] and SwissSideChain databases, allowing predicting peptides with non-canonical amino acids (NCAAs), post-translational modifications (PTMs), D-amino acids, modifications of the peptide terminus and cyclic peptides up to 7-25 residues [20]. Nevertheless, the number of possible modifications was restricted up to 147 NCAAs and 32 PTMs, making it quite restrictive for application due to the growing chemical diversity of peptides.

The emergence of deep learning (DL) has marked a new direction for protein and peptide structure prediction. The AlphaFold (AF) model showed that neural networks can learn distance distributions between residues based on evolution and predict correct protein structures [21,22]; AlphaFold2 (AF2) utilized the same principle in the context of multiple sequence alignments (MSA) and structural templates to produce structures close to experimental ones for CAA proteins [23]. The complementary approach was presented by the ESMFold (EF) method that employed a huge protein language model to obtain evolution and structure information from individual protein sequences without MSA and thereby predicted structures significantly faster without loss of precision [24]. However, all these methods are trained mainly on proteins composed of CAAs and are not reliable for peptide structures with other residues. AF3 is an advanced model with a diffusion-based architecture (atom-level) that demonstrated a substantial improvement in accuracy for predicting biomolecular complexes that include proteins, nucleic acids, small molecules, ions, and modified residues. The introduction of AF3 has significantly improved the usefulness of DL in structural biology and molecular design. However, the current implementation provides support for only a predefined set of 23 PTMs [25]. Therefore, the peptides that have new or uncommon chemical modifications will not be properly accounted for, so it is important to also include the physics-based improvement alongside DL methods.

In addition to the universal predictors described above, several specific frameworks were designed to address unique challenges associated with cyclic and chemically modified peptides. APPTEST utilized a combination of distance and dihedral angle restraints generated based on neural networks and the simulated annealing method to predict the structure of cyclic and linear natural peptides of moderate length [26]. AfCycDesign utilized AF2-based models and included cyclic positional encoding that allowed accurate structure prediction, design and de novo generation of cyclic peptide scaffolds with CAAs [27]. HighFold, an AF-derived model, used special encodings that allowed the consideration of head-to-tail cyclization and disulfide bonding during the structure prediction, which allowed accurate modeling not only of cyclic peptides but also cyclic peptide-protein complexes. Thus, the DL methods could be applied to the important class of therapeutic peptides [28]. Continuing this work, HighFold2 predicted the structure of cyclic peptides containing NCAAs using atom-level residue representation, cyclic positional encoding and structure refinement, while HighFold3 used AF3 models for improving predictions of both linear and cyclic peptides containing NCAAs [29,30]. NCPepFold was specifically designed for cyclic peptides containing NCAAs using cyclic positional encoding and multigranular residue- and atom-level representation [31]. Despite these advances, many existing methods remain focused on specific classes of cyclic or modified peptides and do not provide broad support for the diverse range of chemical modifications encountered in modern peptide therapeutics.

The scope of biomolecular modeling methods has been improved greatly too. RoseTTAFold broadened its All-Atom extended DL-based modeling to the assemblies including proteins, nucleic acids, small molecules, metals, and covalent modifications [32]. OpenFold presented a trainable open-source reimplementation of AF2 [33] and OpenFold3 extended its range of use to proteins, ligands, nucleic acids and antibody complexes [34]. BOLTZ-1 reached AF3 accuracy within an open-source environment for biomolecular complex prediction [35]. HighRelax introduced a physics-based refinement protocol with expanded AMBER parameters covering 139 NCAAs to improve stereochemistry, chirality, and structural quality of AI-predicted models. However, HighRelax functions as a refinement-only tool that depends on AF3-generated structures as input, inheriting AF3’s limitation of supporting only 23 predefined PTMs at the structure generation stage. Consequently, peptides containing modifications outside this predefined set cannot be adequately modelled regardless of the refinement quality applied downstream [36]. While these frameworks represent major advances in biomolecular modelling, they are primarily designed for large proteins or biomolecular complexes. Despite partial support for chemical modifications in some cases, they do not provide the systematic and comprehensive modification coverage required for short chemically modified peptide structure prediction.

To address these challenges, we present PEPstrMOD2, an updated computational framework of PEPstrMOD for predicting the tertiary structures of chemically modified peptides of any length range. PEPstrMOD2 expands the range of chemical coverage of peptide modeling through newly developed libraries of modified residues. These libraries were generated using a standardized AMBER parameterization workflow involving Antechamber, prepgen, parmchk2, and tLEaP to produce force-field compatible parameters for modified residues [37,38]. In addition, PEPstrMOD2 uses MAP input format that enables flexible encoding of chemical modifications directly within peptide sequences [39]. The prediction pipeline integrates DL structure predictors with molecular mechanics refinement, where initial structures generated by AF2 or EF are subsequently refined through energy minimization and short MD simulations. Together, these developments enable PEPstrMOD2 to provide a practical, scalable, and automated framework for accurate prediction of structures of chemically modified peptides while overcoming the length restrictions and limited modification coverage of PEPstrMOD.

## Methods

### Dataset

In order to fully evaluate the ability of PEPstrMOD2 in predicting cyclic and chemically modified peptides, the performance of PEPstrMOD2 was tested using three benchmark databases (Figure 1). The first dataset, named AfCyc, includes 80 cyclic peptides collected from AfCycDesign, with 17 head-to-tail (N-C) cyclic peptides and 63 disulfide-cyclized peptides [27]. The second dataset, ModPep433, includes 433 chemically modified peptides collected from the ModPep dataset that has been used for the evaluation of PEPstrMOD. Details about the construction of the dataset can be found in PEPstrMOD work [20]. Since only 433 out of 501 sequences can be processed by AF3, the common subset was chosen to conduct a fair comparison between all the methods. The third dataset, ModPep16, comprises 16 peptides extracted from the ModPep dataset based on the presence of at least 60% regular secondary structure content (helix + strand) and was used for additional structural evaluation as described previously [20].

**Figure 1.**
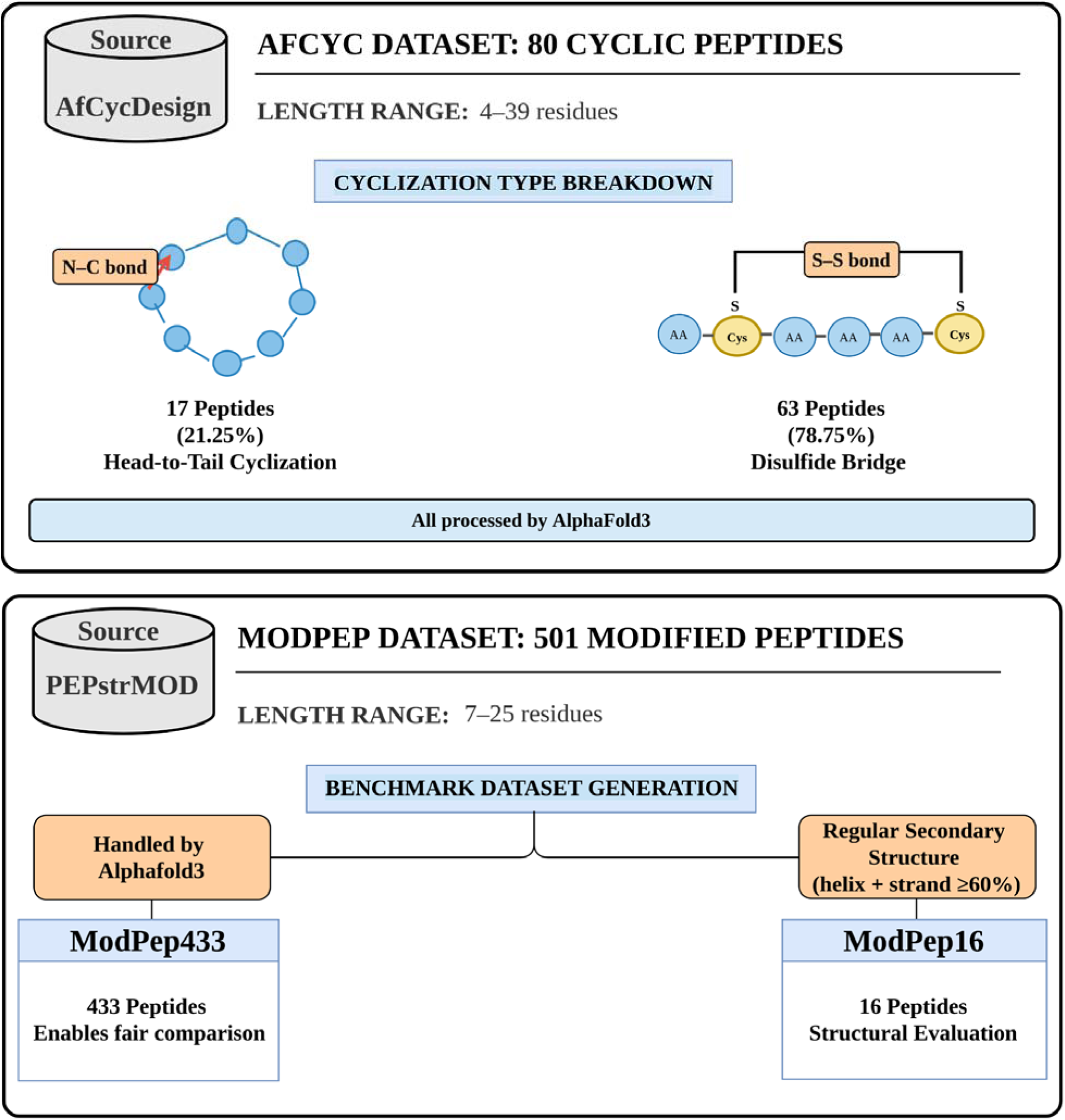
Benchmark datasets used for evaluating PEPstrMOD2. Three datasets were used in this study: AfCyc (80 cyclic peptides), ModPep433 (433 modified peptides), and ModPep16 (16 modified peptides with ≥60% regular secondary structure content).

### Performance measure

To assess the performance of PEPstrMOD2 relative to existing approaches, we compared its predictions with those generated by established peptide structure prediction methods using the same evaluation metrics. For the AfCyc dataset, the predictions from PEPstrMOD2 have been benchmarked with respect to the predicted results obtained by the use of AF3 and AfCycDesign which is the protocol designed for the design and evaluation of cyclic peptides. The modified peptide data sets were benchmarked in respect to their structures predicted using PEPstrMOD and AF3. RMSD values were used to measure the similarity between the predicted structures and experimentally determined peptide structures.

Three types of RMSD were computed: all-atom (AA) RMSD, backbone (BB) RMSD, and C-alpha (CA) RMSD. RMSD calculations were performed using PyMOL. It first superimposes the predicted and experimentally determined structures, and then it calculates the RMSD values. To avoid bias from differences in hydrogen placement, hydrogen atoms were removed from both the predicted and experimental structures before calculating RMSD. For structures determined by NMR containing multiple models, the representative model specified in the Protein Data Bank (PDB) was used as the reference structure. In the absence of any representative model, the first model in the set of NMR structures was taken for the analysis. Distances between sulfur atoms of cysteine residues forming disulfide bonds in cyclic peptides were checked to see if they had preserved correct disulfide bonding in the predicted structures.

For the ModPep16 dataset, secondary-structure preservation was analyzed by using Define Secondary Structure of Proteins (DSSP) assignments [40]. Agreement between experimentally determined and predicted secondary structures was assessed using both the eight-state (Q8: 3_10_helix (G), α-helix (H), π-helix (I), β-stand (E), bridge (B), turn (T), bend (S), and others (C)) and three-state (Q3: helix (H), strand (E), and coil (C)) DSSP classifications [41].

For the ModPep433 dataset, additional structural quality assessments were performed. The local accuracy of the structure was measured through the use of Local Distance Difference Test (LDDT) [42], and the secondary structures assigned by DSSP. The stereochemistry quality was assessed by MolProbity which evaluates steric clash, Ramachandran statistics, rotamer outliers, C-beta deviation, and the covalent geometry of the structure [43]. Together, these metrics provide complementary measures of structural accuracy, secondary-structure preservation, and stereochemical quality.

### Overview of PEPstrMOD

The workflow of the original PEPstrMOD method has been described previously [26690490]. Briefly, PEPstrMOD used predicted secondary structure and β-turn type torsional restraints to make an initial peptide structure. Then, it used AMBER or GROMACS to do energy minimization and short MD simulations. Although this method made it possible to model peptides of length range 7-25 amino acids with different kinds of modifications, it relied on restraint-based structure generation and had limited coverage of chemical modifications.

### PEPstrMOD2 Prediction Pipeline

PEPstrMOD2 automates the process of predicting the three-dimensional structures of chemically modified peptides of any length range (Figure 2). The pipeline starts with peptide sequences that are encoded in the MAP format. This format lets us show various chemical modifications (PTMs, NCAAs, terminal modifications, D-amino acids, and cyclic constraints) right in the peptide sequence. The initial step uses AF2 or EF. These DL models use the input sequence to predict how the peptide will fold and give a basic canonical structural model. Following the prediction of initial structure, the chemical modifications delineated in the MAP input are integrated to the peptide structure. This is accomplished using AMBER-compatible modification libraries designed for PEPstrMOD2. Using the tLEaP module of AMBER, the modified residues are added to the predicted peptide structure while maintaining the overall backbone conformation obtained from the DL models. The peptide structure obtained after the modification process will be subjected to energy minimization using the main engine in AMBER called sander/Particle Mesh Ewald Molecular Dynamics (PMEMD). Minimization of energy will serve to remove any steric clashes that exist in the structure and enhance the local geometry of the peptide structure. Subsequent short MD simulations will be carried out using sander/PMEMD. The refined structure obtained after minimization and MD simulation is reported as the final predicted tertiary structure of the peptide. Overall, our tool offers an automated and scalable framework for predicting peptide structures of different lengths with various chemical modifications, enabled by the integration of MAP-based sequence annotation, DL-based structure prediction, modification incorporation, and molecular mechanics refinement.

**Figure 2:**
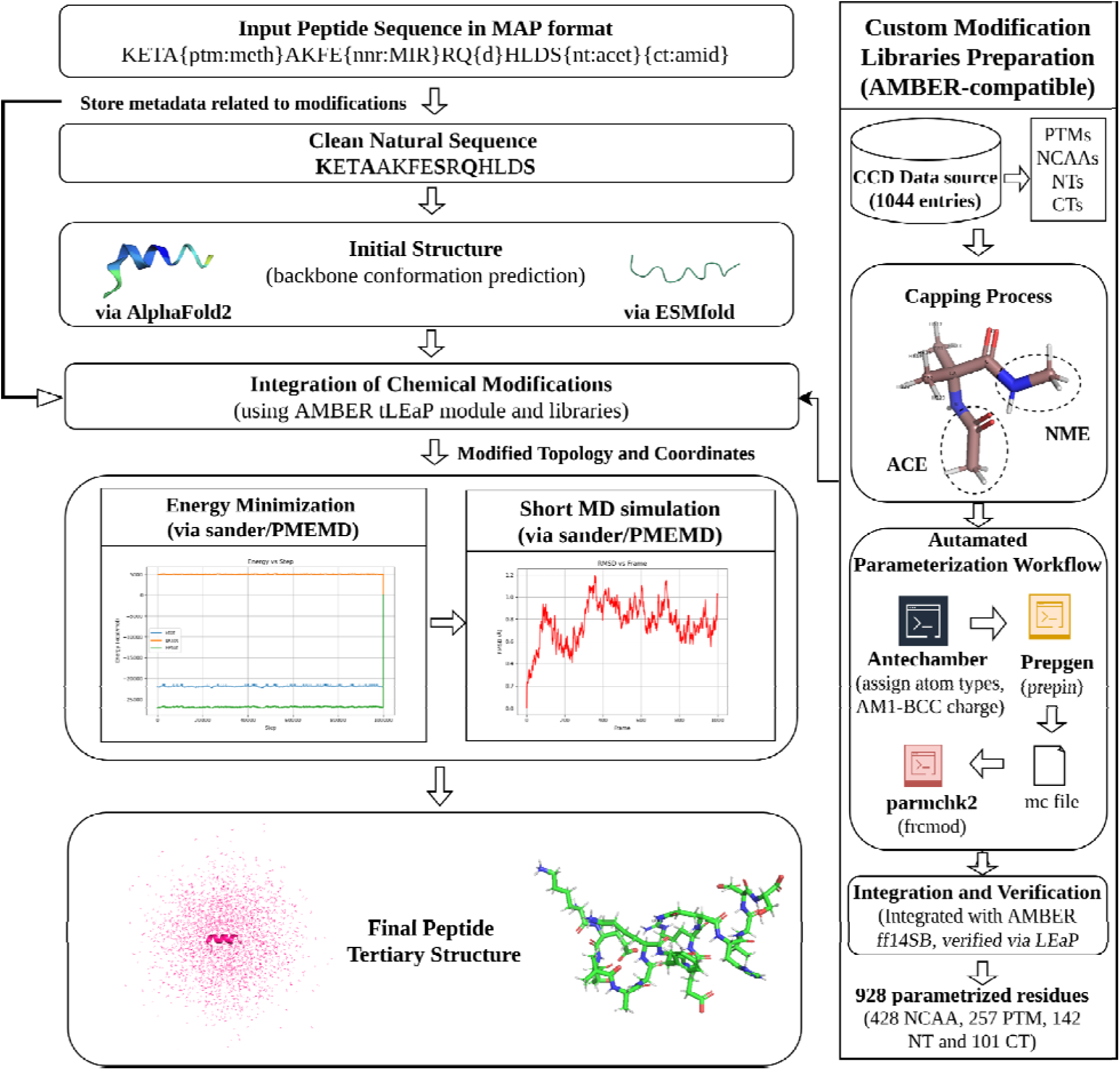
Schematic diagram illustrating the algorithmic procedure followed by PEPstrMOD2.

### Preparation of Modification Libraries

We created expanded libraries of residues that work with the AMBER force-field so that chemically modified peptides can be modeled correctly (Figure 2). We got modified residues from the CCD of the PDB [44] by choosing entries that were marked as L-peptide residues [45]. The residues collected were categorized into four categories, namely NCAAs, PTMs, NTs and CTs. The process used both CCD annotations and manual inspection. In total, 1044 candidate residues were initially identified. All residues were processed through an automated parameterization workflow. The molecular structures were derived from the CCD CIF files and used in the creation of the force-field. In case of modifications that are internal (PTMs and NCAAs), both ends of the molecules were acylated using the ACE (Acetyl) group and NME (N-Methyl Amide) group respectively. For terminal modifications, only the opposite terminus was capped (ACE for CTs and NME for NTs). This ensured that each residue was parameterized in a chemically consistent peptide-like environment. Force-field parameters were generated using the AMBER parameterization pipeline (AMBERTOOLS25). Antechamber was used to assign atom types and compute partial atomic charges using the AM1-BCC charge model. Residue topology files (PREPIN) were generated using prepgen together with custom main-chain (mc) definition files specifying head, tail, backbone atoms, and atoms to be removed during residue construction. Missing parameters were subsequently identified using parmchk2, which generated the corresponding FRCMOD files. The generated parameters were integrated with the AMBER ff14SB protein force-field, and finally tLEaP was used to validate the libraries [37,38]. Generation of the topologies of peptides in tLEaP indicated that the force-field parameters generated were perfectly compatible with the AMBER force-field system.

A total of 928 residues have been parameterized in this way, creating suitable PREPIN and FRCMOD libraries. Other residues in the CCD database have not been converted because of unavailable atoms like boron and other heavy atoms or inadequate structural definition. The library set included in the PEPstrMOD2 software includes 428 NCAAs, 257 PTMs, and 243 termini modifications, including 142 NT and 101 CT residues. This library is helpful for automated addition of chemical modifications while predicting peptide structures.

### Molecular Dynamics Refinement

After the incorporation of the modified residues, the structures of the peptides predicted were refined using energy minimization and short MD simulation to refine local geometry and remove steric clashes. All simulations were performed using the AMBER24 simulation package with parameters compatible with the PEPstrMOD2 modification libraries. Initially, energy minimization was carried out for 2000 steps using a combination of steepest descent (1000 steps) and conjugate gradient algorithms (1000 steps) to optimize the peptide structure. A non-bonded cutoff of 10 Å was used. Next, the system was heated for 50 picoseconds (ps) using NVT ensemble (at 300 K temperature) followed by equilibration for 50 ps using NPT ensemble (at 1 bar pressure). The equilibrated system was subsequently subjected to short MD simulations at 300 K and 1 bar (timestep = 1 femtosecond) to further relax the structure and stabilize the conformation in the presence of chemical modifications. In a vacuum environment, the system was heated for 50 ps at 300 K followed by production MD at 300 K with no periodic boundary conditions. For performing simulation in water, TIP3P water model was used and for simulations in a hydrophobic environment, a methane box was used. The simulation time can be selected by the user in ps or nanoseconds (ns). The refined structure obtained after minimization and MD refinement was reported as the final predicted tertiary structure of the peptide. Simulations can be performed either in vacuum, water, or a hydrophobic environment. All the parameters corresponding to MD simulations in AMBER used in PEPstrMOD2 are provided in Additional File Table A1.

## Results and Discussion

To evaluate the performance of PEPstrMOD2 on different sets of peptide modifications and structures, we benchmarked the algorithm on several data sets. For each peptide, the structure was predicted using PEPstrMOD2 and compared with the experimentally determined structures available in the PDB. The final predicted structure after MD simulation was taken as the Last Frame (LF) of the trajectory. In addition, to assess structural consistency, we also computed representative structures based on the highest populated cluster and the average structure of all frames. The cluster containing the highest number of trajectory frames was selected as the highest populated cluster. The centroid structure corresponding to this cluster was used as the representative structure. The average structure was generated by aligning all frames in PyMOL followed by coordinate averaging using custom Python scripts. AA-RMSD, BB-RMSD, and CA-RMSD were calculated for all three sets of structures (LF, cluster representative, and average structure) by superimposing them with their corresponding experimental structures.

### Performance of PEPstrMOD2 on AfCyc Dataset

To test the performance of PEPstrMOD2 on cyclic peptides, we employed the AfCyc data set, which includes 80 experimentally validated cyclic peptide structures, 17 peptides being head-to-tail cyclized (N-C) and 63 peptides disulfide bonded (S-S). The performance of several prediction models, including PEPstrMOD2, PEPstrMOD, AfCycDesign, AF3, AF2, and EF, was assessed based on AA-, BB-, and CA-RMSD scores. To analyze the influence of MD refinement conditions, the simulations were conducted in the vacuum and hydrophilic environments for 100 ps and 1 ns time steps.

Under 100 ps vacuum MD simulation conditions, the best-performing refinement model was the PEPstrMOD2_AF2 average model that scored an average RMSD of 2.21 Å (AA), 1.63 Å (BB), and 1.67 Å (CA). The results of 100 ps vacuum MD simulations are shown in Supplementary Table S1. An extension of 100 ps vacuum MD simulations up to 1 ns did not significantly change the RMSD scores, namely 2.31 Å (AA), 1.70 Å (BB), and 1.75 Å (CA). Detailed results for the 1 ns vacuum simulations are provided in Supplementary Table S2.

When simulations were performed in an explicit hydrophilic environment, improved structural agreement with experimentally determined structures was observed. Under 100 ps hydrophilic simulations, the PEPstrMOD2_AF2 average model achieved the best overall refinement performance with average RMSD values of 2.05 Å (AA), 1.50 Å (BB), and 1.51 Å (CA). Detailed results for the 100 ps hydrophilic simulations are provided in Supplementary Table S3. Increasing the simulation time to 1 ns produced only modest additional changes, resulting in RMSD values of 2.11 Å (AA), 1.52 Å (BB), and 1.60 Å (CA). Detailed results for the 1 ns hydrophilic simulations are provided in Supplementary Table S4. These observations suggest that most structural relaxation occurs during the early stages of MD refinement, while longer simulations provide limited additional benefit. The enhanced performance in the hydrophilic simulations is possibly caused by the presence of solvent molecules that stabilize peptide structures further and decrease the probability of steric clash during refinement. This leads to better agreement with the experimental structure of cyclic peptides compared to the vacuum simulation.

Among the DL-based approaches, AF3 demonstrated the best results with the lowest RMSD values on all criteria, reaching the average RMSD values of 1.13 Å (AA), 0.87 Å (BB), and 0.90 Å (CA). AfCycDesign demonstrated a good performance level with RMSD values of 1.82 Å (AA), 1.33 Å (BB), and 1.35 Å (CA). AF2 provided average RMSD values of 2.41 Å (AA), 1.88 Å (BB), and 2.09 Å (CA) and was inferior in terms of accuracy to EF, which had RMSD values of 3.02 Å (AA), 2.20 Å (BB), and 2.61 Å (CA).

PEPstrMOD2 remained competitive with AF3, performed better than EF and AF2 after refinement, and was almost consistent with the structures generated by AfCycDesign (Table 2). This shows that the combination of DL-based starting model and physics based molecular mechanics refinement greatly enhances the cyclic peptide structure prediction while maintaining the chemically plausible conformation. Moreover, PEPstrMOD2 is capable of accommodating a wider range of cyclic and chemically modified peptides without needing any special training for cyclic peptides.

**Table 2.** Comparison of PEPstrMOD2 average structure performance on AfCyc dataset with different methods.

|  | AA |  |  | BB |  |  | CA |  |  |
| --- | --- | --- | --- | --- | --- | --- | --- | --- | --- |
|  | Init | Min | Sim | Init | Min | Sim | Init | Min | Sim |
| **PEPstrMOD2_EF (Avg)** | 2.81 | 2.42 | 2.45 | 2.10 | 1.81 | 1.80 | 2.43 | 1.90 | 1.87 |
| **PEPstrMOD2_AF2 (Avg)** | 2.38 | 2.15 | 2.05 | 1.82 | 1.59 | 1.50 | 2.06 | 1.59 | 1.51 |
| **AfCycDesign** | 1.82 | - | - | 1.33 | - | - | 1.35 | - | - |
| **EF** | 3.02 | - | - | 2.20 | - | - | 2.61 | - | - |
| **AF2** | 2.41 | - | - | 1.88 | - | - | 2.09 | - | - |
| **AF3** | 1.13 | - | - | 0.87 | - | - | 0.90 | - | - |

*AA: All Atoms; BB: Backbone Atoms; CA: C-alpha atoms; Init: Initial structure without simulation; Min: Minimized Structure; Sim: Simulated structure; EF: ESMFold; Avg: Average model; AF2: Alphafold2; AF3: Alphafold3*

To further evaluate whether the predicted structures preserved correct disulfide connectivity, we analyzed the distances between sulfur atoms of cysteine residues forming disulfide bridges (in 63 peptides having S-S bond) using structures refined under 100 ps vacuum simulation conditions. The usual disulfide bond distance found for experimentally determined structures is about 2.0 Å. This was true for all the refined structures produced through PEPstrMOD2 modeling, with a range of ∼1.9-2.1 Å observed for the S-S distance, suggesting that chemically valid disulfide bond geometry had been retained. In contrast, structures generated directly by some DL-based models occasionally produced incorrect cysteine pairing geometries, resulting in larger S-S distances. The refinement stage implemented in PEPstrMOD2 corrected these geometries and restored physically realistic disulfide bond lengths during molecular mechanics optimization. Detailed disulfide bond distance analyses are provided in Supplementary Table S5.

### Performance of PEPstrMOD2 on ModPep Dataset

To evaluate the performance of PEPstrMOD2 on peptides containing diverse chemical modifications, we used the ModPep dataset, which consists of 501 experimentally determined modified peptide structures obtained from the PDB and previously used in the original PEPstrMOD study. The performance of different prediction methods was evaluated using AA-, BB-, and CA-RMSD metrics.

For this analysis, structure prediction was performed using AlphaFold2, followed by 100 ps MD simulations in an explicit hydrophilic environment using the TIP3P water model for structural refinement. Owing to the large-scale computational requirements associated with MD simulations for 501 peptides, the final frame, i.e., LF obtained after simulation was used for structural evaluation. Cluster-based representative structures and average structure calculations were not employed because of their substantially higher computational power at this scale. The performance of PEPstrMOD2 was compared to PEPstrMOD which is summarized in Supplementary Table S6. For PEPstrMOD, the initial predicted structures showed average RMSD values of 5.12 Å (AA), 3.96 Å (BB), and 4.21 Å (CA). After energy minimization, the RMSD values improved slightly to 5.01 Å (AA), 3.85 Å (BB), and 4.12 Å (CA).

In contrast, PEPstrMOD2 produced substantially improved initial models with average RMSD values of 3.58 Å (AA), 2.69 Å (BB), and 2.97 Å (CA). Following energy minimization, the structures showed RMSD values of 4.15 Å (AA), 3.26 Å (BB), and 3.08 Å (CA). The minimized structures were subsequently subjected to 100 ps MD simulations in hydrophilic conditions, resulting in RMSD values of 4.43 Å (AA), 3.36 Å (BB), and 3.26 Å (CA).

Such results demonstrate that PEPstrMOD2 delivers significantly better initial structural predictions in comparison with PEPstrMOD, owing to the use of the DL structure prediction approach. Although the RMSD values increase slightly after energy minimization and MD simulations, these refinement steps play an important role in relaxing the predicted structures, removing steric clashes, and ensuring chemically realistic geometries for modified residues.

Due to the large size of the ModPep dataset, extended MD simulations such as vacuum simulations or longer simulation times (1 ns) were not performed on this 501 dataset. More extensive analyses examining the effects of simulation environment and simulation time were performed on the ModPep16 dataset with well-defined regular secondary structure (>60%), which are described in the following sections.

#### Performance of PEPstrMOD2 on ModPep433 Dataset

To further examine the performance comparison between PEPstrMOD2 and AF3, we performed the analysis of the peptides that were successfully predicted by AF3. Due to limitations in handling certain chemically modified residues, AF3 was able to process only 433 sequences from the ModPep 501 dataset. Therefore, comparisons between AF3, PEPstrMOD, and PEPstrMOD2 were performed on this common subset of 433 peptides (Supplementary Table S7).

On this subset, AF3 achieved average RMSD values of 4.67 Å (AA), 3.65 Å (BB), and 4.04 Å (CA). In comparison, PEPstrMOD produced RMSD values of 5.00 Å (AA), 3.87 Å (BB), and 4.12 Å (CA) for the initial structures and 4.88 Å (AA), 3.74 Å (BB), and 4.03 Å (CA) after energy minimization. The initial RMSD value of PEPstrMOD2 was 3.56 Å (AA), 2.68 Å (BB), and 2.98 Å (CA). After the energy minimization, the RMSD was 4.18 Å (AA), 3.31 Å (BB), and 3.07 Å (CA), and after 100 ps MD simulations, the RMSD was 4.49 Å (AA), 3.43 Å (BB), and 3.28 Å (CA).

All in all, it can be concluded that PEPstrMOD2 greatly improves the prediction of chemically modified peptides by providing better starting structures via DL-based prediction while keeping the strong refinement pipeline, based on physics.

Next, the ModPep433 dataset was divided into four groups depending on peptide lengths (7-10, 11-15, 16-20, and 21-25 amino acids) to examine the effect of peptide size on the prediction quality (Supplementary Table S8). Performance of AF3, PEPstrMOD minimized model, and PEPstrMOD2_AF2 (LF model) at different peptide sizes is shown in Table 3. Overall, PEPstrMOD2_AF2 consistently achieved lower RMSD values than both AF3 and PEPstrMOD across most length categories. PEPstrMOD2_AF2 produced the most accurate predictions for the 7-10 residue peptides (average RMSD = 3.44 Å (AA), 2.31 Å (BB), and 2.55 Å (CA)). RMSD increased with increasing length of peptides within 11-15 and 16-20 residue subcategories due to increased conformational flexibility of longer peptides.

**Table 3.**
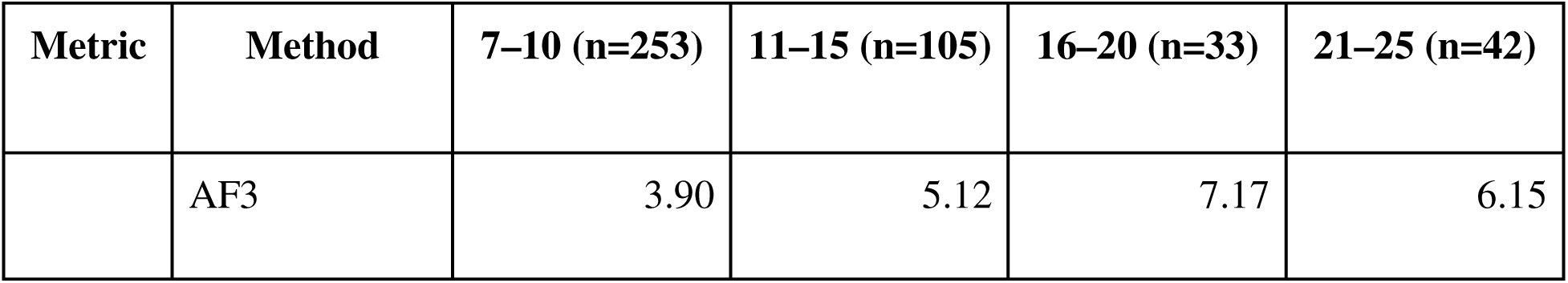
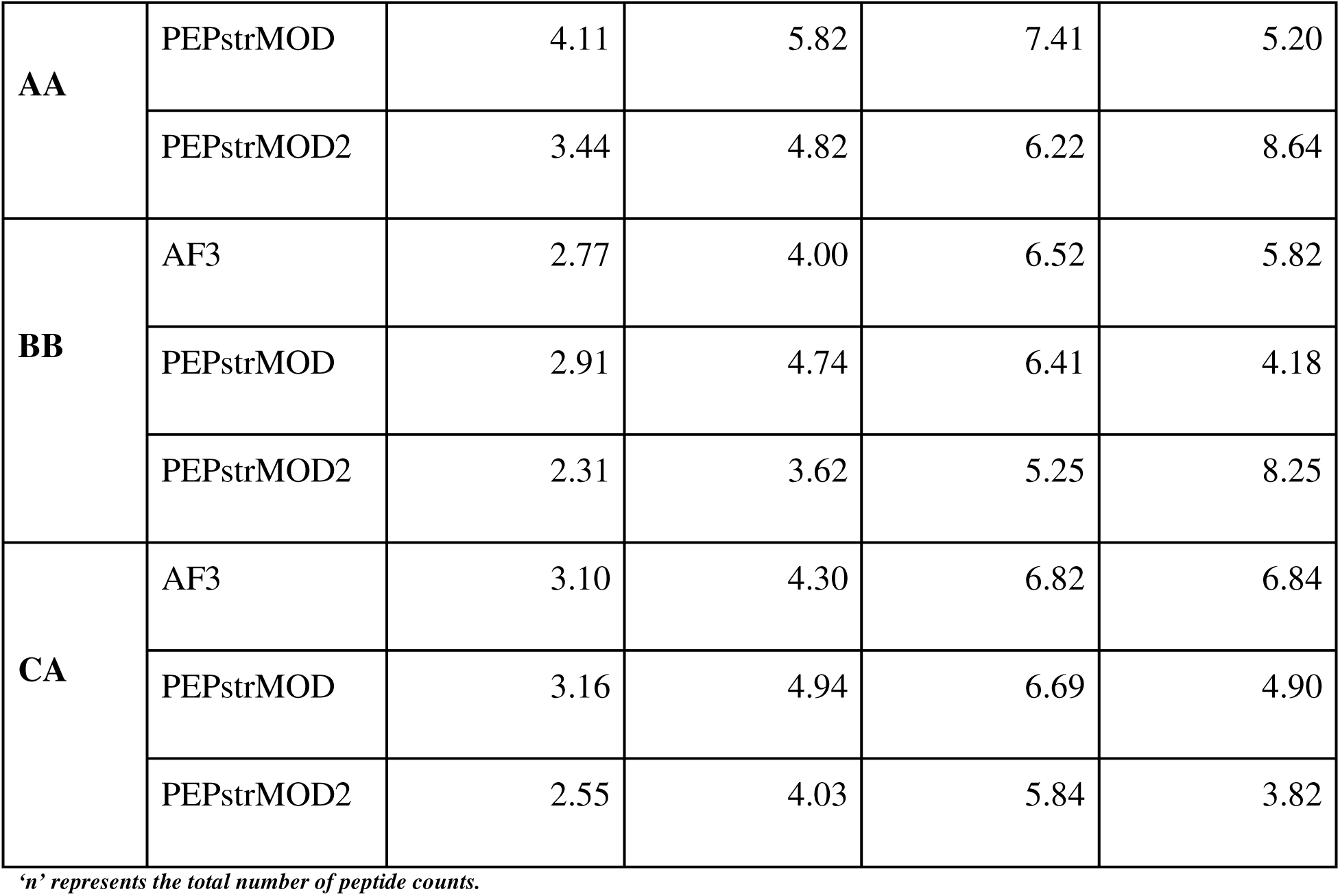
Performance comparison of different methods on peptides with different length distributions.

When analyzing the longest peptides (21-25 residues), we observed that correlation between peptide length and structural prediction accuracy was more complicated. PEPstrMOD2_AF2 obtained relatively high AA- and BB-RMSD values (8.64 Å and 8.25 Å, correspondingly), but produced the lowest CA-RMSD among all tested predictors (3.82 Å). Further analysis of this subcategory showed that it was mostly composed of collagen-like peptides with repetitive sequence and triple-helical structure. Such structures are highly sensitive to small rotational, translational, or register shifts between peptide chains. Consequently, the overall trajectory of the peptide backbone can remain largely preserved, resulting in relatively low CA-RMSD values, while local differences in backbone geometry, hydrogen-bonding patterns, and chain packing produce substantially larger deviations at the AA and BB levels. As a result, several collagen-like peptides showed good agreement with the experimental structures in terms of overall fold but comparatively higher AA- and BB-RMSD values. Therefore, the observed discrepancy between atomistic and CA-level accuracy in this subset likely reflects the structural characteristics of collagen-like peptides rather than a simple dependence on peptide length.

Thus, we can conclude that peptide length influences accuracy of predictions in a nonmonotonous way and depends on a number of factors such as peptide length, conformational flexibility, and structural class. The enrichment of collagen-like peptides in the longest length category highlights the importance of considering conformational flexibility, and sequence-specific structural features, in addition to sequence length, when evaluating peptide structure prediction methods.

Further structural quality assessments were carried out on structures generated through AF3 and PEPstrMOD2_AF2 (LF model, hydrophilic environment, 100 ps). These analyses included evaluation of secondary structure agreement using DSSP, stereochemical quality assessment using the MolProbity server, and local structural accuracy using LDDT.

Secondary structure agreement between predicted and experimentally determined peptide structures was evaluated using DSSP. Secondary structure assignments were analyzed using both Q8 and Q3 DSSP classification, and the results are summarized in Supplementary Table S9. For ModPep433, average Q8 and Q3 agreements for PEPstrMOD2_AF2 were 62.44% and 75.37%, respectively, versus 49.87% and 69.87% for AF3. However, these numbers need to be considered carefully because the majority of peptides in ModPep433 are short, very flexible and coil-rich, with few regular elements of secondary structure. In other words, DSSP agreement depends mainly on the ability to correctly recognize coils but not on prediction of helical or β-strand structures. Therefore, DSSP was used as an auxiliary measure of structure similarity while RMSD and LDDT were utilized as key metrics of the prediction quality.

Local accuracy of the predicted structures was measured using the LDDT metric. Calculation of LDDT score was feasible for 430 peptides out of 433 peptides from the ModPep433 dataset. Three peptides (3SEM_C, 3ZS2_D, and 4JXT_B) were excluded due to the presence of difference in amino acid composition or peptide length in reference and predicted structures that prevented direct LDDT comparison. Distribution of LDDT scores for AF3 and PEPstrMOD2_AF2 are presented in Table 4 and detailed LDDT scores for each individual peptide are shown in Supplementary Table S10.

**Table 4.** Distribution of LDDT scores for peptide structures predicted by AF3 and PEPstrMOD2_AF2 on the ModPep433 dataset.

| LDDT Range | AF3 Count | AF3 % | PEPstrMOD2_A<br>F2 Count | PEPstrMOD2_A<br>F2 % |
| --- | --- | --- | --- | --- |
| $\geq 90$ | 43 | 10.00 | 39 | 9.07 |
| 80-89 | 29 | 6.74 | 124 | 28.84 |
| 70-79 | 56 | 13.03 | 113 | 26.28 |
| 60-69 | 82 | 19.07 | 80 | 18.60 |
| 50-59 | 115 | 26.74 | 49 | 11.40 |
| <50 | 105 | 24.42 | 25 | 5.81 |
| **Total** | **430** | **100** | **430** | **100** |

*Interpretation of LDDT score ranges is adapted from AlphaFold confidence score guidelines [46] and reflects the expected local structural accuracy of predicted models. $\geq 90$ : backbone and side chains predicted with very high accuracy; 70–89: correct backbone with minor side-chain deviations; 60–69: partial backbone correctness; 50–59: backbone largely incorrect; <50: poor structural agreement with very low accuracy.*

The PEPstrMOD2_AF2 model showed a better average LDDT value (0.73) than AF3 (0.62), implying an increased backbone precision and structural similarity with experimentally determined peptides. In addition, 64.19% of predicted peptides structures in the PEPstrMOD2_AF2 model obtained LDDT values ≥0.70, while only 29.77% structures were predicted by the AF3 model, reflecting an increase in the percentage of locally accurate peptides after refinement. Figure 3 illustrates that PEPstrMOD2_AF2 outperforms the AF3 model at most of the LDDT cutoff values, especially at ≥0.70 and ≥0.80. Also, there was a noticeable decrease in the percentage of peptides with low LDDT values (LDDT<0.50) from 24.42% in AF3 to 5.81% in the PEPstrMOD2_AF2 model. All of these results reflect the improvement in the structural similarity between residues of the predicted peptide structure and the experimentally determined one after MD refinement.

**Figure 3.**
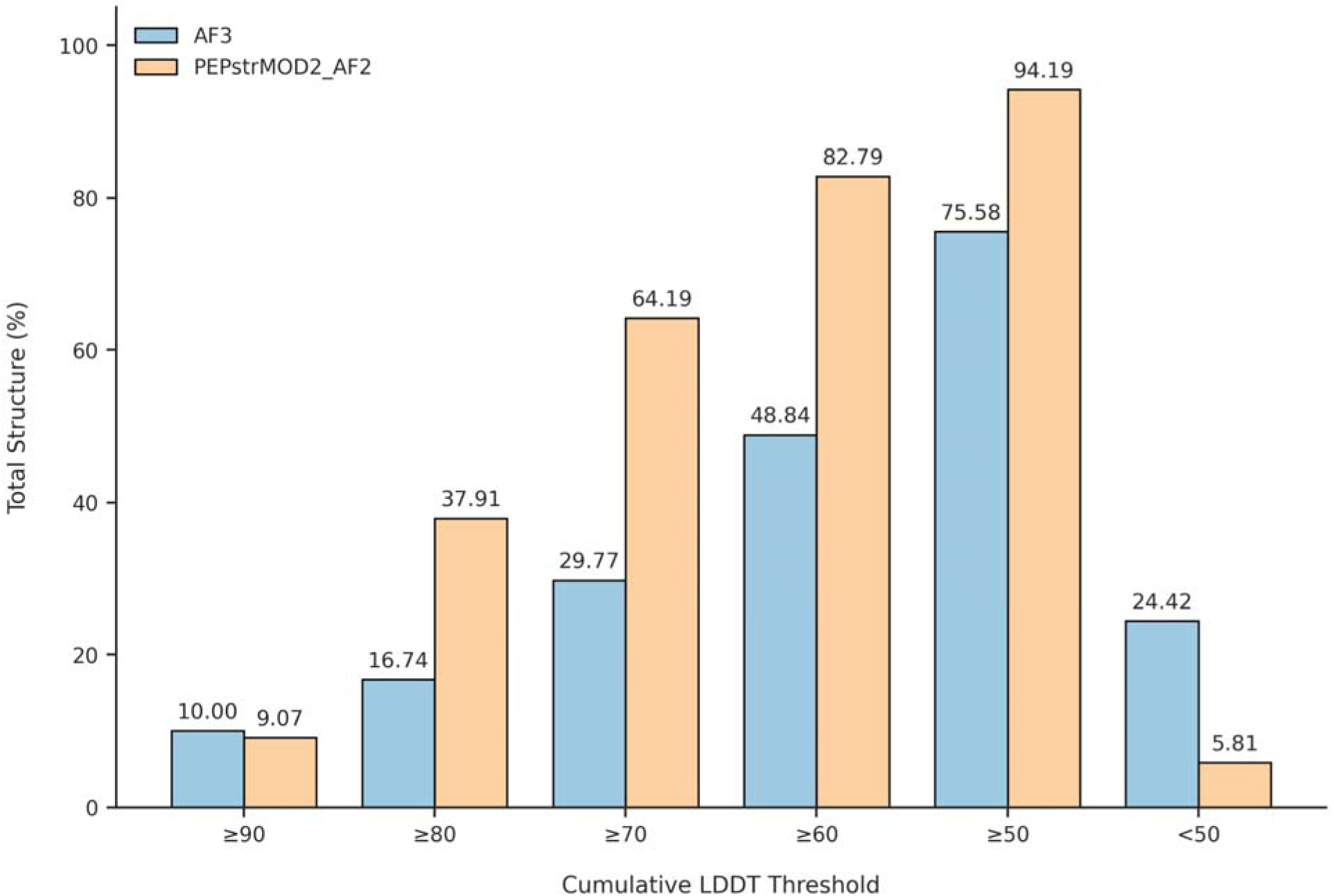
Cumulative percentage of peptide structures exceeding different LDDT cutoffs for AF3 and PEPstrMOD2_AF2 models on ModPep433 dataset. PEPstrMOD2_AF2 outperform AF3 at most LDDT cutoffs, reflecting the improved local structural accuracy after MD refinement

To further quantify the stereochemical quality of peptide structures, all 433 experimental structures (Original), AF3 predictions, and refined PEPstrMOD2_AF2 models were evaluated using MolProbity. Summary statistics are shown in Table 5. The experimental structures had overall good stereochemical quality, with a MolProbity score of 1.84 ± 1.20 and a clashscore of 12.49 ± 23.47. On the other hand, stereochemical quality of the AF3 predictions was poor, with higher clashscores (176.97 ± 124.64), higher Ramachandran outliers (7.59 ± 13.86%), and high MolProbity score (3.95 ± 1.12). This shows that although AF3 produces peptide structures with global plausibility, problems with local steric clashes and backbone conformation errors persist. Structures predicted using PEPstrMOD2_AF2 refinement pipeline had very improved stereochemical quality. This is seen in the low clashscore of 0.34 ± 1.55, which is approximately 500 times lower than the AF3 predictions. Ramachandran outliers decreased from 7.59 ± 13.86% to 1.03 ± 5.33%, while rotamer outliers were reduced from 30.41 ± 20.68% to 2.12 ± 5.40%. Moreover, MolProbity score improved from 3.95 ± 1.12 for AF3 predictions to 0.94 ± 0.54 after refinement.

**Table 5.** Comparison of MolProbity stereochemical validation metrics (Mean±Standard Deviation) for experimental structures (Original), AF3 predictions, and refined (100ps, hydrophilic) PEPstrMOD2_AF2 LF model in the ModPep433 dataset.

| **Metric** | **Original** | **AF3** | **PEPstrMOD2_AF2** | **Recommended values** |
| --- | --- | --- | --- | --- |
| **Clashscore** | 12.49±23.47 | 176.97±124.64 | 0.34±1.55 | <10 |
| **Ramachandran outliers (%)** | 4.78±12.40 | 7.59±13.86 | 1.03±5.33 | <0.05 |
| **Ramachandran favored (%)** | 84.82±24.80 | 77.72±26.22 | 90.01±18.89 | >98 |
| **MolProbity score** | 1.84±1.20 | 3.95±1.12 | 0.94±0.54 | Lower is better |
| **Rotamer Outliers (%)** | 9.46±14.48 | 30.41±20.68 | 2.12±5.40 | <0.3 |

*Values represent mean ± standard deviation calculated across 433 peptide structures. MolProbity scores and clashscores suggest better stereochemical quality. The suggested thresholds are taken according to the MolProbity validation recommendations [47] and are mainly based on globular proteins; hence, direct comparisons should be done with caution in case of short peptides and structurally restricted peptide systems.*

Despite the percentage of residues in favorable Ramachandran regions being lower than those usually reported for globular protein structures with high resolution, there was no significant difference between the three datasets. This result was anticipated since peptide datasets contain many short, glycine-rich, proline-rich, and structurally restricted peptides that often occupy unfavorable backbone regions.

Other validation metrics, such as Cβ deviation, RMS bond deviation, and RMS angle deviation, displayed the same pattern and validated the increased stereochemical quality of the refined PEPstrMOD2_AF2 structures. Complete values of all validation metrics are provided in Supplementary Table S11. Distribution of MolProbity score and Clashscore in 433 peptide structures is presented in Figure 4. As can be seen from the graph of empirical cumulative distribution function (ECDF), there is a strong tendency of the PEPstrMOD2_AF2 structures towards the lower values of MolProbity score and Clashscore in comparison with the AF3 and experimental structures (Figure 4A and 4C). In line with this, the violin plot demonstrates that PEPstrMOD2_AF2 structures have the lowest values of median MolProbity score and Clashscore and the compact distribution while the AF3 structures have significantly higher values of both scores and more variance (Figure 4B and 4D). Together, these results indicate that stereochemical improvements are observed across the majority of peptides rather than being driven by a small number of outlier structures.

**Figure 4.**
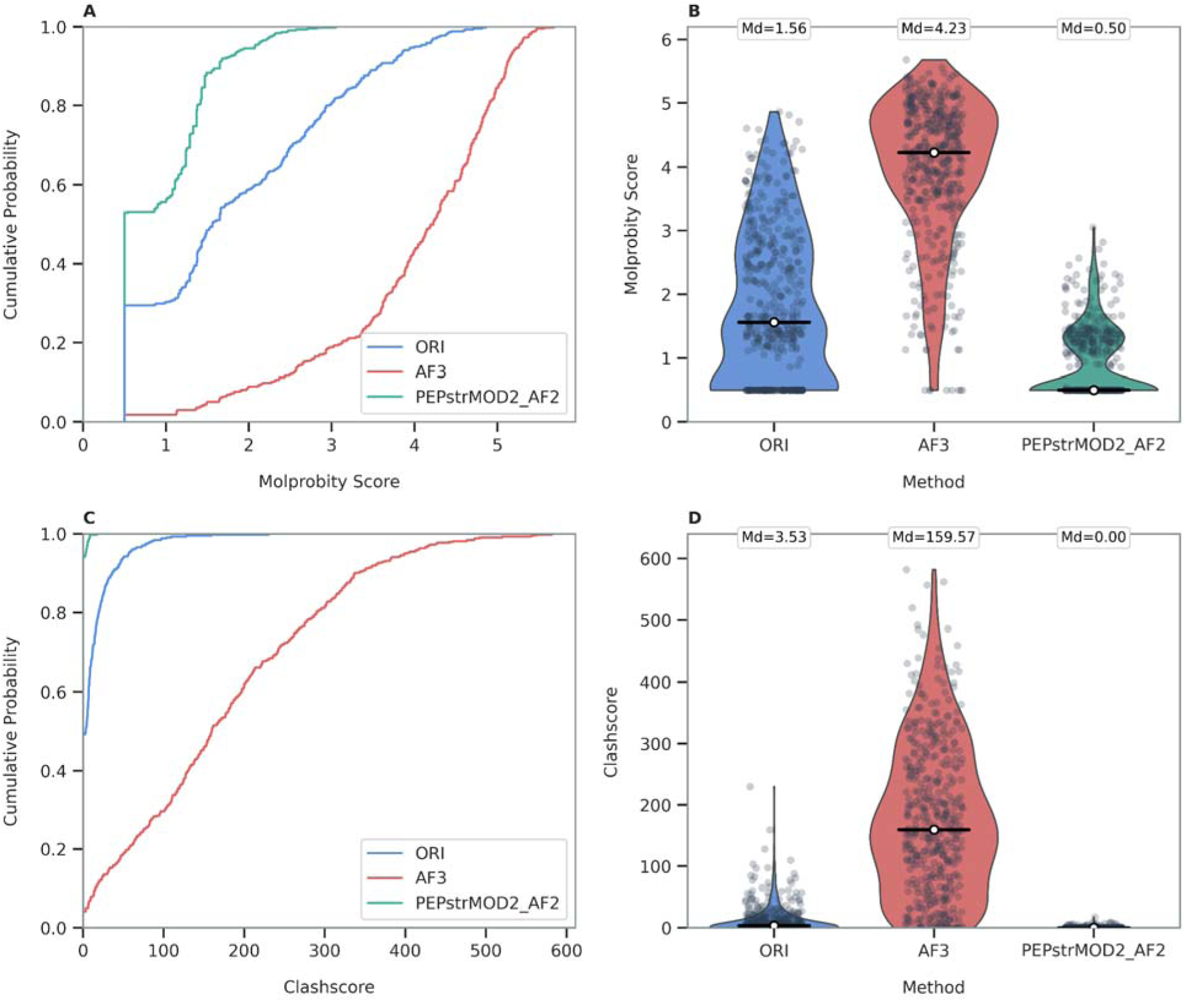
Distribution of MolProbity scores and Clashscores for ModPep433. (A, C) ECDF of MolProbity scores and Clashscores of experimental structures (ORI), AF3 models, and PEPstrMOD2_AF2 refined structures (n = 433 per dataset). (B, D) Violin plots of score distributions. White circles show median (Md) values.

Although the PEPstrMOD2_AF2 refinements gave better MolProbity scores compared to the experimental structures, this is due to higher geometric ideality rather than biological accuracy, as energy minimization brings the structures closer to optimized bond lengths, bond angles, torsionals and non-bonded contacts, while the experimentally determined structures could still have minor geometric imperfections caused by experimental error, thermal motion, crystal packing, or biologically meaningful conformational strain. In summary, it can be concluded that the PEPstrMOD2 structure refinement pipeline significantly enhances peptide stereochemistry by reducing clashes and optimizing backbone and side-chain geometries.

#### Performance of PEPstrMOD2 on ModPep16 Dataset

The ModPep16 dataset contains 16 peptides and represents a subset of the ModPep dataset comprising peptides with regular secondary structure content (helix and strand) ≥60%, as defined in the original PEPstrMOD study. To evaluate the performance of PEPstrMOD2, peptide structures were first predicted using EF and AF2 and subsequently refined using the PEPstrMOD2 pipeline involving energy minimization and short MD simulations. Predicted structures were compared with experimentally determined structures using RMSD after structural superposition. Their performance was compared with PEPstrMOD only. We did not include AF3 in this comparison because it was able to process only 2 out of the 16 sequences due to limitations in handling certain chemically modified residues present in the dataset. Therefore, AF3 results were not considered for systematic benchmarking on the ModPep16 dataset. With respect to the 100 ps vacuum simulations, the PEPstrMOD2_EF average structure model exhibited the lowest RMSD values of 3.75 Å (AA), 2.86 Å (BB) and 3.27 Å (CA), a considerable enhancement when compared to the performance of the original PEPstrMOD framework which yielded RMSD values of 5.46 Å (AA), 4.05 Å (BB) and 4.31 Å (CA). Detailed results for the 100 ps vacuum simulations are provided in Supplementary Table S12. Extending the vacuum simulations to 1 ns did not result in any significant improvements. Detailed results for the 1000 ps vacuum simulations are provided in Supplementary Table S13.

In order to determine the impact of solvent conditions on structural refinements, the PEPstrMOD2 models were also subjected to explicit hydrophilic simulations using the TIP3P water model. With regard to the 100 ps hydrophilic simulations, the PEPstrMOD2_AF2 average structure model obtained RMSD values of 2.75 Å (AA), 1.89 Å (BB) and 2.20 Å (CA), which are more refined structures than their counterparts from vacuum simulations. Results of the 100 ps hydrophilic simulations are shown in Supplementary Table S14. The best performances were found in the 1000 ps hydrophilic simulations, whereby the PEPstrMOD2_AF2 average structure model attained RMSD values of 2.50 Å (AA), 1.75 Å (BB) and 2.07 Å (CA). These results are summarized in Table 6, while detailed results for the 1000 ps hydrophilic simulations are provided in Supplementary Table S15.

**Table 6.** The performance of different methods on peptides in the ModPep16 dataset (AA-RMSD). All our models were subjected to 1000 ps MD simulations in a hydrophilic environment.

| PDB_ID | Length | PEPstrMOD_sim | PEPstrMOD2_EF | PEPstrMOD2_AF2 |
| --- | --- | --- | --- | --- |
| 1fevA | 15 | 4.06 | 3.04 | 1.54 |
| 1rbdS | 15 | 5.13 | 0.47 | 1.24 |
| 1tkqB | 15 | 6.73 | 6.72 | 6.55 |
| 1z3lS | 15 | 5.20 | 0.85 | 2.27 |
| 1z3mS | 15 | 6.50 | 2.04 | 1.5 |
| 1z3pS | 15 | 5.83 | 1.86 | 1.97 |
| 2ap8A | 20 | 2.72 | 1.01 | 1.01 |
| 2dprA | 21 | 3.07 | 1.53 | 1.81 |
| 2fx8P | 12 | 6.06 | 1.61 | 1.67 |
| 2k7lB | 19 | 8.50 | 3.68 | 1.75 |
| 2rlnS | 15 | 6.25 | 1.91 | 1.12 |
| 3cmhA | 15 | 7.92 | 6.09 | 3.44 |
| 3kmzC | 19 | 2.30 | 0.75 | 0.85 |
| 3zs2D | 25 | 7.39 | 2.14 | 2.24 |
| 4lkaB | 12 | 5.95 | 1.09 | 5.87 |
| 6cmhA | 21 | 5.79 | 7.23 | 5.17 |
| **Average** |  | **5.59** | **2.63** | **2.50** |

A few peptides showed relatively large RMSD values across multiple prediction methods. According to structural analysis, the cases described above fall under those peptides which have experimentally derived structures highly dependent on the presence of stabilization factors like oligomeric interfaces, ligand interactions, or particular solvents. For example, 1TKQ_B corresponds to a β-helix gramicidin structure stabilized by inter-chain hydrogen bonds and Cs binding, 3ZS2_D represents a β-strand fragment from the insulin dimer interface, 4LKA_B is a histone H3 tail peptide that adopts a helical conformation only upon protein binding, and 6CMH_A is an endothelin-1 analogue with flexible terminal regions reported in NMR studies. Without the presence of such factors, when simulated in isolation, conformational flexibility is increased and higher RMSD values are acquired.

To investigate the effect of these structures, RMSD calculations were repeated after excluding these four structures (Table 7). Removal of outliers led to a substantial improvement in structural agreement in both PEPstrMOD and PEPstrMOD2. In particular, the AA-RMSD of the PEPstrMOD2_AF2 average model improved to 1.68 Å. Similar improvements were observed for BB- and CA-RMSD values. Detailed results are reported in Supplementary Table S16.

**Table 7.** The performance of different methods on peptides in the ModPep12 dataset (AA-RMSD)

| **PDB_ID** | **Length** | **PEPstrMOD** | **PEPstrMOD2_EF** | **PEPstrMOD2_AF2** |
| --- | --- | --- | --- | --- |
| 1fevA | 15 | 4.06 | 3.04 | 1.54 |
| 1rbdS | 15 | 5.13 | 0.47 | 1.24 |
| 1z3lS | 15 | 5.20 | 0.85 | 2.27 |
| 1z3mS | 15 | 6.50 | 2.04 | 1.5 |
| 1z3pS | 15 | 5.83 | 1.86 | 1.97 |
| 2ap8A | 20 | 2.72 | 1.01 | 1.01 |
| 2dprA | 21 | 3.07 | 1.53 | 1.81 |
| 2fx8P | 12 | 6.06 | 1.61 | 1.67 |
| 2k7lB | 19 | 8.50 | 3.68 | 1.75 |
| 2rlnS | 15 | 6.25 | 1.91 | 1.12 |
| 3cmhA | 15 | 7.92 | 6.09 | 3.44 |
| 3kmzC | 19 | 2.30 | 0.75 | 0.85 |
| **Average** |  | **5.30** | **2.07** | **1.68** |

To see the effect of terminal residues, which are known to exhibit high conformational variability in both experimental and simulated (PEPstrMOD2_AF2 average model, 1000 ps, hydrophilic) structures, we removed them. There was a substantial improvement in structural agreement (Supplementary Table S16). In particular, the AA-RMSD of PEPstrMOD2_AF2 improved to 1.15 Å (from 1.68 Å), while the corresponding value for experimentally derived structures (simulated at the same condition) was 0.93 Å. A detailed comparison is shown in the Additional File Figure A1, Table A2. These results indicate that a significant portion of the observed structural deviation arises from flexible terminal segments rather than inaccuracies in the predicted core structure.

Because the ModPep16 dataset was specifically selected for peptides containing at least 60% regular secondary structure (helix and strand), secondary-structure preservation was also evaluated using DSSP assignments. Comparison of DSSP states between experimentally determined and PEPstrMOD2_AF2 average model (1000ps, hydrophilic) yielded average agreement scores of 69.26% for the Q8 classification and 79.35% for the Q3 classification (Supplementary Table S17). Since there is higher agreement between DSSP for Q3 class, it shows that the overall structure class (helix, strand, or coil) remains almost intact even after prediction and refinement, although discrepancies in detailed DSSP assignment could be because of local structural features such as turns, bends, or helices of different subtypes. However, some peptides had lower agreement, especially those in which the conformation of the peptide relies upon the presence of stabilizing forces from other sources such as interactions between proteins or ligand binding or certain types of solvent. For instance, peptides such as 4LKA_B and 1TKQ_B showed low DSSP agreement because of their flexibility or dependence on the context.

In summary, it has been shown that the PEPstrMOD2 refinement pipeline considerably improves the stereochemical quality of peptide structures. The combination of DL method for prediction and energy minimization leads to generation of precise peptide structures with low number of steric clashes and precise backbone conformation.

## Software Availability

PEPstrMOD2 is available as a web server (https://webs.iiitd.edu.in/raghava/pepstrmod/) and also as a standalone version through GitHub (https://github.com/raghavagps/PEPstrMOD2), along with Docker support, for easier installation and scalable predictions of peptide structures. PEPstrMOD2 can be installed locally through a standalone package.

## Webserver Implementation and User Interface Workflows

In order to illustrate the operation of the PEPstrMOD2 workflow, we outline an example of running a prediction task for a modified peptide (Figure 5). The user submits their peptide sequence (up to 100 residues) via the online interface in the form of the MAP inline notation with modifications. The user can use a number of templates for D-residues, PTMs, non-standard residues, cyclization, disulfide bonds, and terminal cap residues along with a live parser which parses the submitted sequence and identifies the modifications in real time (Figure 5A). The user chooses the structure prediction approach (ESMFold) and sets up the simulation environment.

**Figure 5.**
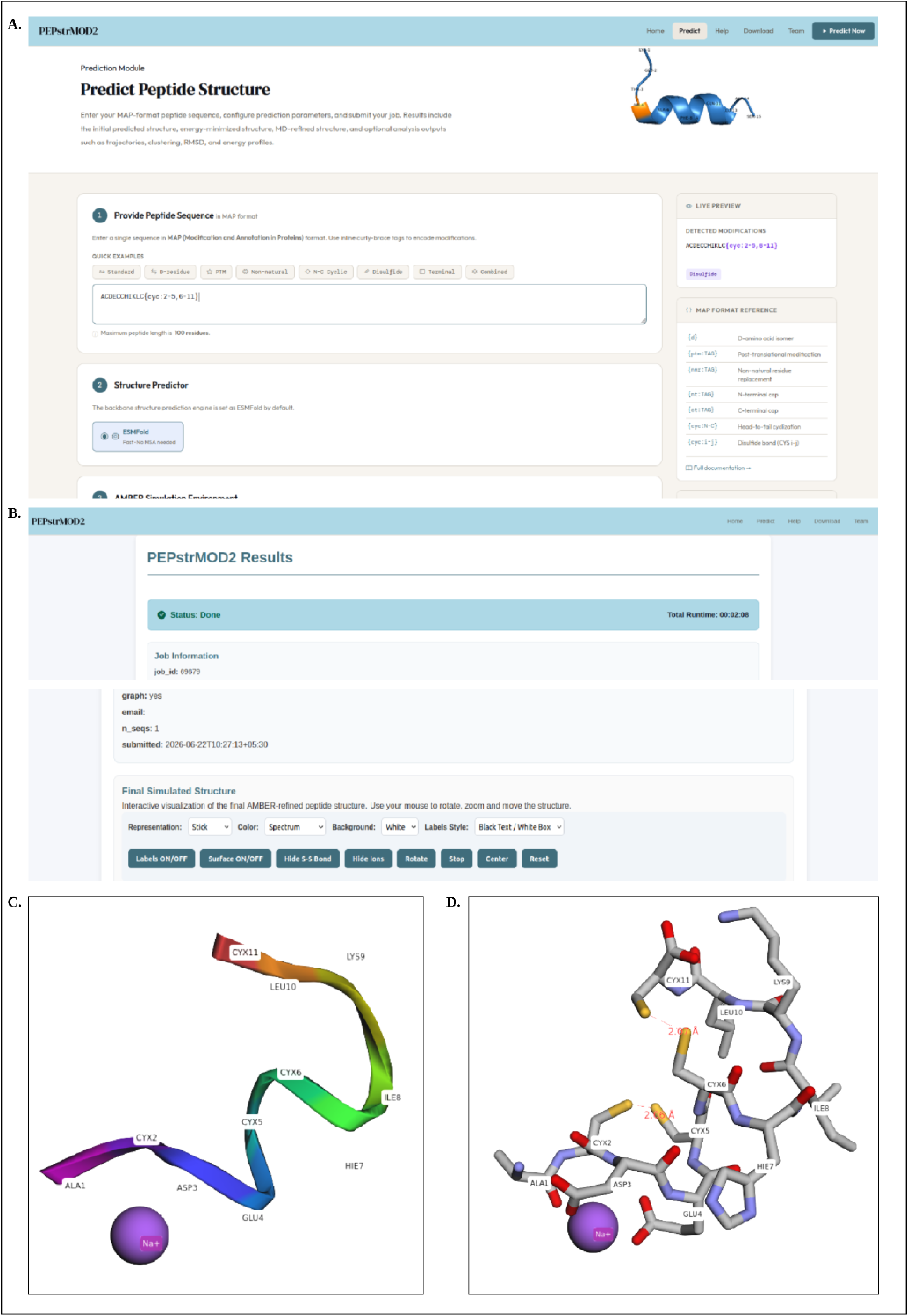
Interface and procedure of the PEPstrMOD2 web server for the disulfide-cyclized peptide ACDECCHIKLC{cyc:2-5,6-11}. (A) Submission form displaying the input sequence in the MAP format and preview of modifications. (B) Results page displaying the status of the computation, computation time, and input parameters. (C) Cartoon depiction of the optimized structure with the Na^+^ ion. (D) Stick depiction of the optimized structure showing two automatically identified disulfide bonds (CYX2–CYX5, CYX6–CYX11) with their measured bond distances.

After the job is completed, the results are delivered to a results page, which provides information about the job status, job runtime, and the entered parameters (Figure 5B). The refined structure is depicted in a browser-based manner through the 3Dmol.js program, with controls of representation style, color mode, background, and labels along with additional controls for surface rendering, disulfide bonds, and ions display. When cartoon representation is chosen, the structure is visualized with a smooth backbone trace and spectrum coloring, and the predicted counterion is visualized as a sphere (Figure 5C). Changing to stick representation provides the same structure in high resolution, while highlighting cysteines forming disulfide bonds and calculating their S-S distance (Figure 5D).

### Execution Time

The time taken to predict the structure of the peptides using PEPstrMOD2 is dependent on various parameters like the length of the peptides, the deep learning method used for prediction, and the computational power. Using DL predictors like AF2 or EF to create the first structure takes a lot of computing power. However, the step involving the generation of the MD is the most time-consuming stage in the procedure, particularly for longer simulations (greater than 1 ns). In contrast, the subsequent refinement steps, including energy minimization and short MD simulations (100 ps), require less computational time. The prediction time depends on the availability of CPU and GPU resources. The availability of webserver and docker support enables reproducible deployment and facilitates large-scale peptide structure prediction workflows.

### Conclusion

Peptide-based therapeutics play an important role in the treatment of numerous diseases and disorders. The knowledge of the structural characteristics of peptides is important to study the biological roles of peptides and to develop peptide-based drugs through rational drug design. In this work, we developed PEPstrMOD2, an updated computational framework for predicting the tertiary structures of chemically modified peptides of variable lengths. The method expands the chemical coverage of peptide modeling through newly developed libraries of NCAAs, PTMs, and terminal modifications and integrates DL-based structure prediction with molecular mechanics refinement. Benchmark evaluation on multiple datasets demonstrates that PEPstrMOD2 can effectively model peptides containing diverse chemical modifications. PEPstrMOD2 is freely available as both a webserver and a standalone package. The standalone implementation of PEPstrMOD2 allows local deployment and integration into automated peptide modeling workflows, enabling large-scale prediction tasks that may not be feasible using server-based DL prediction systems. PEPstrMOD2 is expected to be a valuable tool for researchers who are interested in peptide therapeutics, peptide design, and structural bioinformatics.

### Limitations

Despite the great potential of PEPstrMOD2 to extend the peptide structure prediction to chemically modified peptides, certain limitations remain. The method relies on predefined libraries of modified residues, and therefore peptides containing modifications not included in the current libraries cannot be directly modeled. In addition, the accuracy of the predicted structures partly depends on the quality of the initial models generated by DL predictors such as AF2 or EF. Future developments may focus on expanding the coverage of modification libraries and improving modeling of complex peptide architectures.

## Abbreviations

AA: All Atom
ACE: Acetyl
AF: AlphaFold
AF2: AlphaFold2
AF3: AlphaFold3
BB: Backbone
CA: C-Alpha
CAAs: Canonical Amino Acids
CCD: Chemical Component Dictionary
CT: C-terminal
DL: Deep Learning
DSSP: Define Secondary Structure of Proteins
EF: ESMFold
HMM: Hidden Markov Model
LF: Last Frame
MD: Molecular Dynamics
MSAs: Multiple Sequence Alignments
NCAAs: Non-Canonical Amino Acids
NME: N-Methyamide
NT: N-terminal
Ns: nanoseconds
PDB: Protein Data Bank
PMEMD: Particle Mesh Ewald Molecular Dynamics
PTMs: Post-Translational Modifications
ps: picoseconds
RMSD: Root Mean Square Deviation

## Supplementary Materials

Additional File (.docx): Supplementary methods and analyses, including AMBER simulation parameters (Table A1), comparison of experimentally determined and predicted structures for the ModPep12 dataset (Figure A1), and RMSD analysis of simulated structures relative to experimentally determined structures (Table A2).

Supplementary File (.xlsx): Detailed benchmarking results of PEPstrMOD2 on the AfCyc, ModPep433, and ModPep16 datasets, including MD simulation results under different environments and simulation lengths, disulfide bond analysis, secondary-structure comparisons, LDDT scores, and MolProbity validation statistics (Tables S1–S17).

## Authors’ contributions

SJ collected the modified residues. SJ, NKM, SR, PK prepared custom force-field libraries. SJ implemented the algorithms and developed the prediction pipeline model. Varun, SJ, and NKM collected and curated the datasets. SJ, NKM, SR benchmarked the datasets. SJ, NKM, Varun, GPSR analyzed the results. SJ, NKM created the front-end and back-end of the webserver. SJ, NKM, SR, GPSR penned the manuscript. GPSR coordinated the project. All authors have read and approved the final manuscript.

## Acknowledgements

Authors are thankful to the University Grants Commission (UGC) for fellowships and financial support and the Department of Computational Biology, IIITD New Delhi, for infrastructure and facilities. The authors thank Anshuma Yadav for providing support for MD simulations.

## Data Availability Statement

All the datasets used in this study are available at the “PEPstrMOD2” web server, https://webs.iiitd.edu.in/raghava/pepstrmod2/download.html.

