## Additional File for "PEPstrMOD2: Next-generation tertiary structure prediction of chemically modified and non-natural peptides"

### 1. Department of Computational Biology, Indraprastha Institute of Information Technology, Okhla Phase 3, New Delhi-110020, India.

2. Indian Institute of Science Education and Research (IISER) Pune, Dr Homi Bhabha

Road, Pune, Maharashtra, 411008, India.

**Mailing Address of Authors**

***Corresponding Author**

Prof. Gajendra P. S. Raghava

Department of Computational Biology

Indraprastha Institute of Information Technology, Delhi

New Delhi, India - 110020

	Website: [http://webs.iiitd.edu.in/raghava](http://webs.iiitd.edu.in/raghava/)

**Table A1 AMBER parameters for performing energy minimization and molecular dynamics using AMBER25**

| **Vacuum Environment** | | |
| --- | --- | --- |
| **Energy Minimization** | | imin=1, ntmin=1, maxcyc=2000, ncyc=1000, ntb=0, igb=0, cut=10.0 |
| **Molecular Dynamics (MD)** | **Heating** | imin=0, irest=0, ntx=1, ntb=0, ntc=2, ntf=2, cut=10.0, igb=0, tempi=0, temp0=300, ntt=3,  gamma_ln=2, nstlim=50000, dt=.001,ig=-1, ntpr=100, ntwx=100 |
|  | **Production MD** | imin=0, irest=0, ntx=5, ntb=0, ntc=2, ntf=2, cut=10.0, igb=0, tempi=300, temp0=300, ntt=3,  gamma_ln=2, nstlim=100000, dt=.001,ig=-1, ntpr=100, ntwx=100 |
| **Hydrophilic/Hydrophobic Environment** | | |
| **Energy Minimization** | | imin=1, maxcyc=2000, ntmin=1, ncyc=1000, ntb=1,cut=10.0 |
| **Molecular Dynamics (MD)** | **NVT ensemble** | imin=0,irest=0,ntx=1, ntb=1,ntc=2,ntf=2, cut=10.0, tempi=0,temp0=300,ntt=3,gamma_ln=2,  nstlim=50000,dt=.001,ig=-1, ntpr=100,ntwx=100 |
|  | **NPT ensemble** | imin=0, irest=1, ntx=5,ntr=0, ntb=2, taup=2, pres0=1, ntp=1, ntc=2, ntf=2, cut=10.0,  tempi=300,temp0=300,ntt=3, gamma_ln=2, nstlim=50000, dt=.001,ig=-1, ntpr=100,  ntwx=100 |
|  | **Production MD** | imin=0, irest=1, ntx=5,ntr=0, ntb=2, taup=2, pres0=1, ntp=1, ntc=2, ntf=2, cut=10.0,  tempi=300,temp0=300,ntt=3, gamma_ln=2, nstlim=100000, dt=.001,ig=-1, ntpr=100,  ntwx=100 |

**Figure A1 Comparison of the original simulated structure from PDB and PEPstrMOD2_AF2 (Average Model) with the original PDB structures from PDB for the ModPep12 dataset (after removing the highly flexible terminal regions)**

**1FEV_A**: 4-11

| Original_sim vs Original | Pepstrmod2_af2_sim vs Original |
| --- | --- |
| 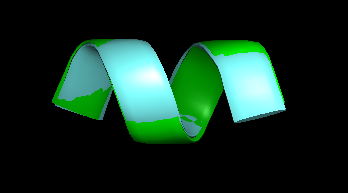 | 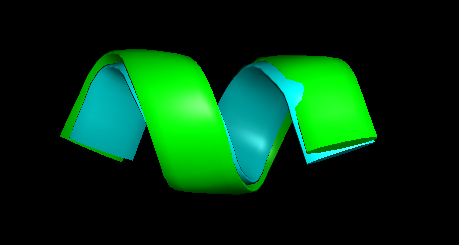 |

**1RBD_S**: 4-11

| Original_sim vs Original | Pepstrmod2_af2 vs Original |
| --- | --- |
| 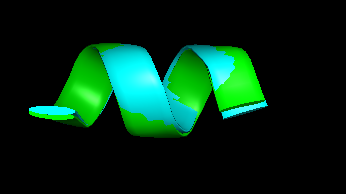 | 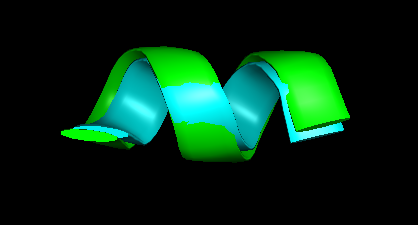 |

**1Z3L_S**: 4-11

| Original_sim vs Original | Pepstrmod2_af2 vs Original |
| --- | --- |
| 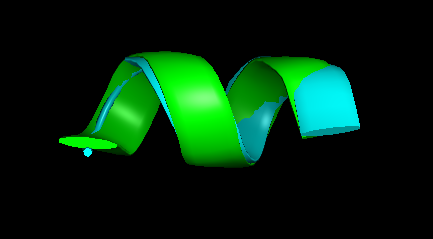 | 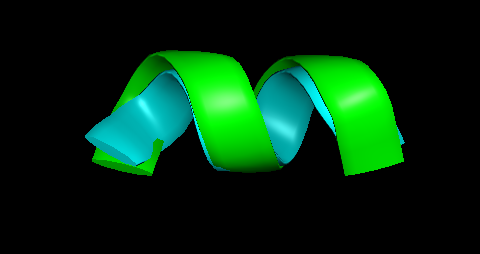 |

**1Z3M_S**: 4-11

| Original_sim vs Original | Pepstrmod2_af2 vs Original |
| --- | --- |
| 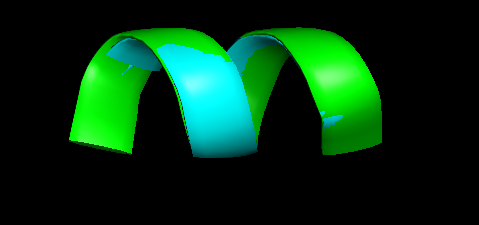 | 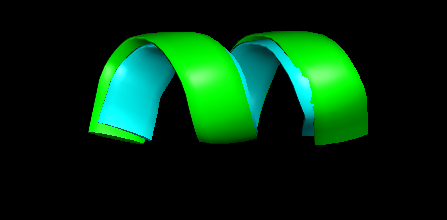 |

**1Z3P_S**: 4-11

| Original_sim vs Original | Pepstrmod2_af2 vs Original |
| --- | --- |
| 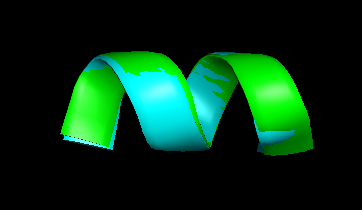 | 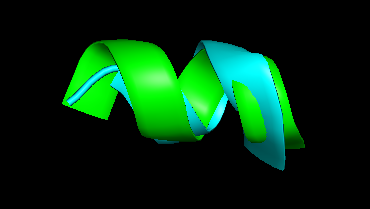 |

**2AP8_A**: 5-18

| Original_sim vs Original | Pepstrmod2_af2 vs Original |
| --- | --- |
| 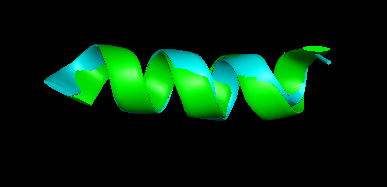 | 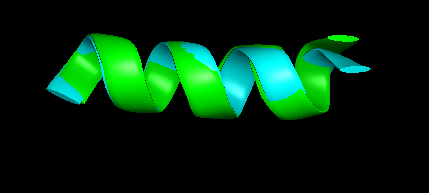 |

**2DPR_A**: 3-18

| Original_sim vs Original | Pepstrmod2_af2 vs Original |
| --- | --- |
| 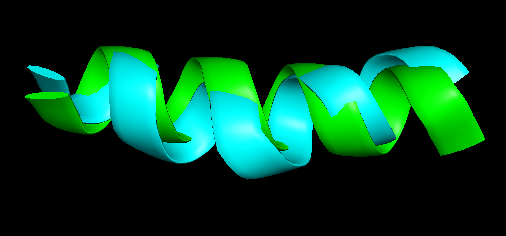 | 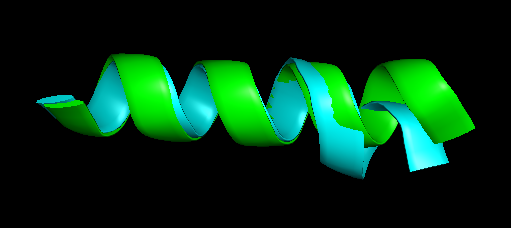 |

**2FX8_P**: 2-10

| Original_sim vs Original | Pepstrmod2_af2 vs Original |
| --- | --- |
| 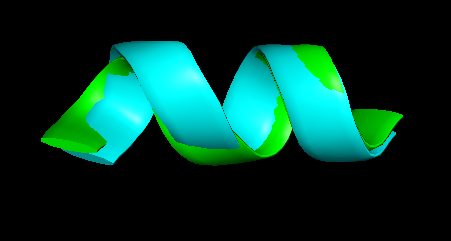 | 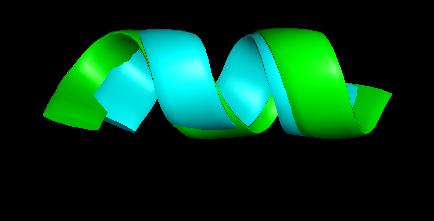 |

**2K7L_B**: **5-16**

| Original_sim vs Original | Pepstrmod2_af2 vs Original |
| --- | --- |
| 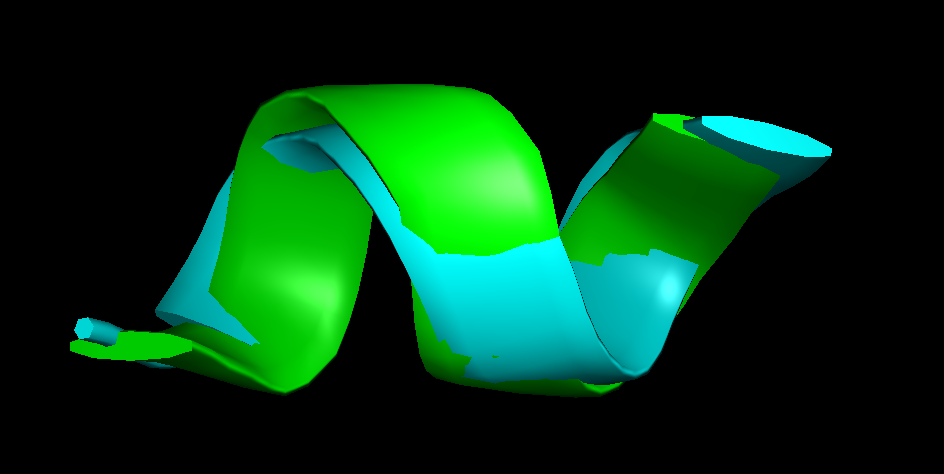 | 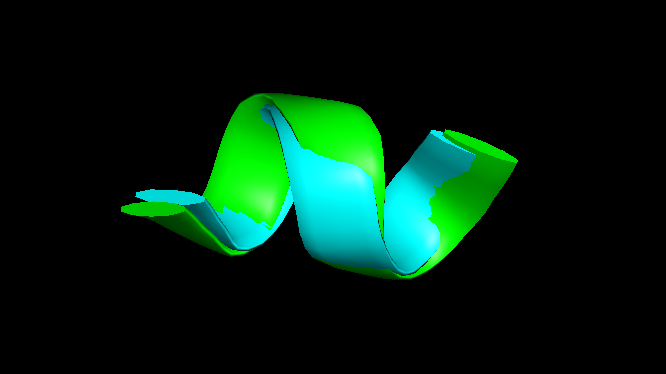 |

**2RLN_S**: 4-11

| Original_sim vs Original | Pepstrmod2_af2 vs Original |
| --- | --- |
| 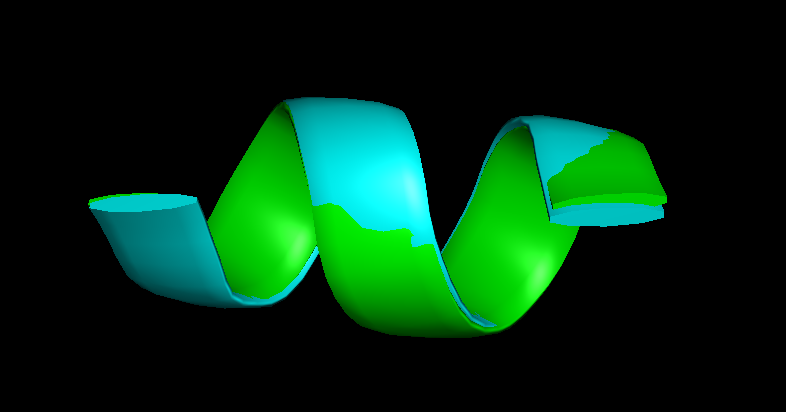 | 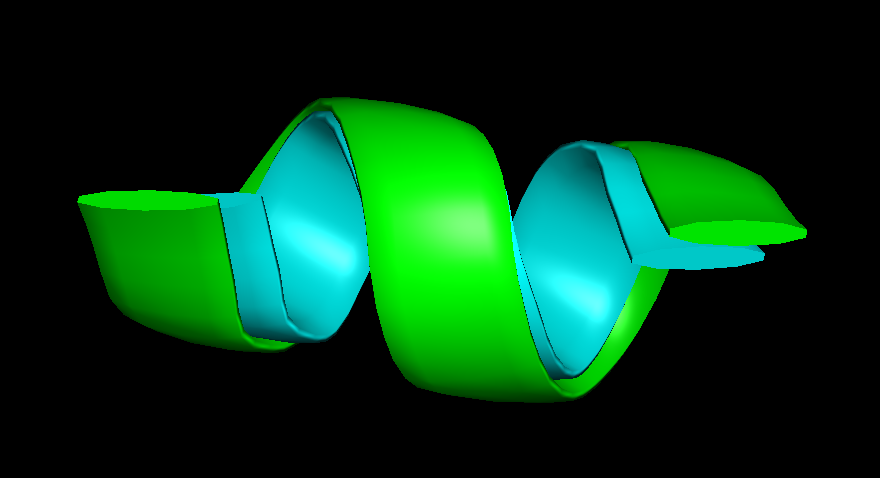 |

**3CMH_A**: 3-11

| Original_sim vs Original | Pepstrmod2_af2 vs Original |
| --- | --- |
| 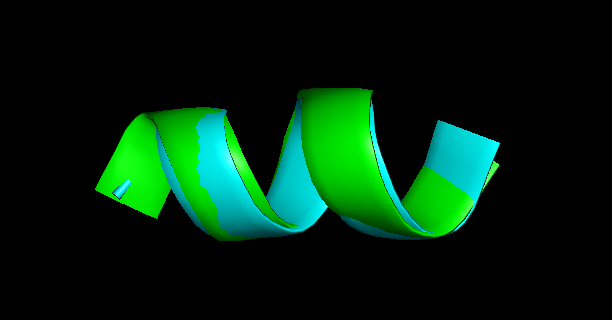 | 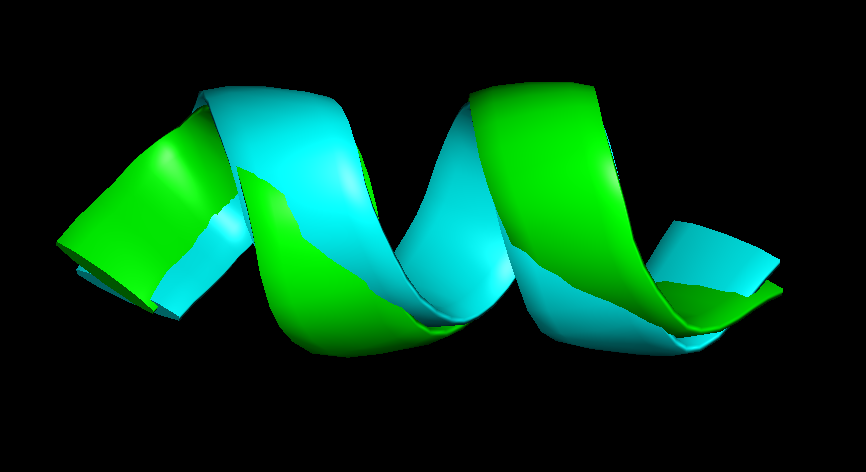 |

**3KMZ_C**: 5-16

| Original_sim vs Original | Pepstrmod2_af2 vs Original |
| --- | --- |
| 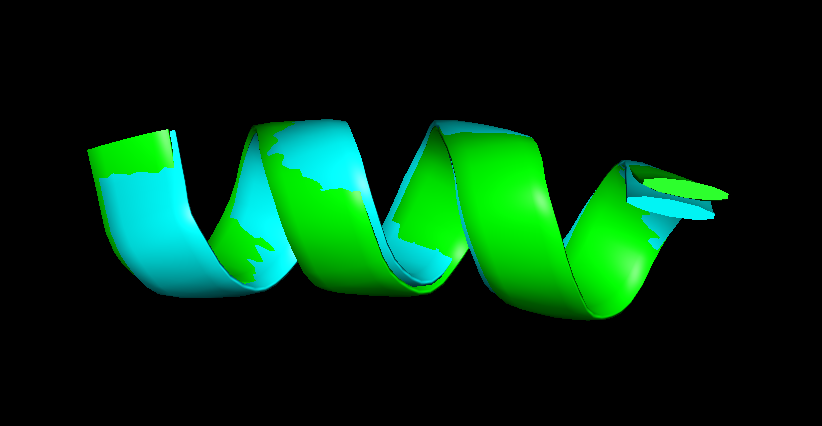 | 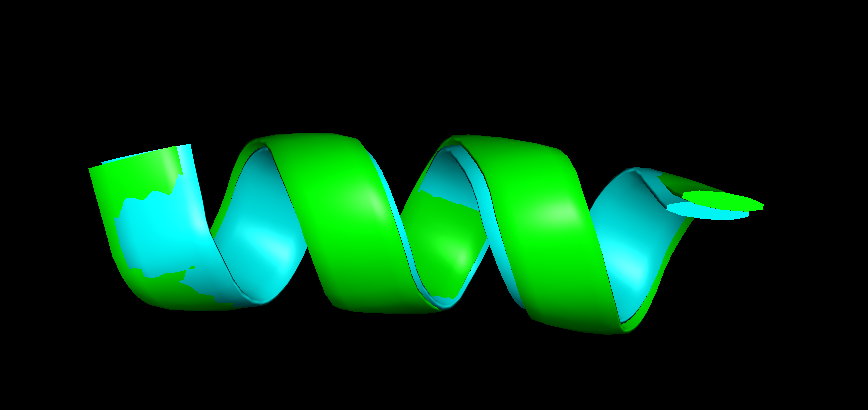 |

Simulated structures are shown in cyan and the original pdb structure in green. Their RMSD values are reported in Table 2.

**Table A2: RMSD values of simulated original structure and PEPstrMOD2_AF2 (Average, 1000ps, hydrophilic) with original PDB structures**

| **Original PDB ID** | **Original_sim** | | | **PEPstrMOD2_AF2** | | |
| --- | --- | --- | --- | --- | --- | --- |
|  | **AA** | **BB** | **CA** | **AA** | **BB** | **CA** |
| 1fevA | 0.46 | 0.27 | 0.26 | 0.63 | 0.33 | 0.34 |
| 1rbdS | 0.37 | 0.29 | 0.31 | 0.96 | 0.48 | 0.54 |
| 1z3lS | 0.83 | 0.74 | 0.63 | 0.84 | 0.62 | 0.65 |
| 1z3mS | 0.34 | 0.25 | 0.23 | 0.51 | 0.44 | 0.46 |
| 1z3pS | 0.61 | 0.35 | 0.35 | 1.93 | 1.60 | 1.64 |
| 2ap8A | 1.21 | 0.57 | 1.11 | 0.60 | 0.47 | 0.57 |
| 2dprA | 2.06 | 1.41 | 1.40 | 1.70 | 1.06 | 0.38 |
| 2fx8P | 1.59 | 0.30 | 0.92 | 1.60 | 1.16 | 1.16 |
| 2k7lB | 1.57 | 0.78 | 0.83 | 1.78 | 0.46 | 0.49 |
| 2rlnS | 0.28 | 0.18 | 0.21 | 1.01 | 0.69 | 0.71 |
| 3cmhA | 1.19 | 0.65 | 0.73 | 1.77 | 0.77 | 0.84 |
| 3kmzC | 0.60 | 0.46 | 0.49 | 0.50 | 0.37 | 0.40 |
| **Average** | **0.93** | **0.52** | **0.62** | **1.15** | **0.70** | **0.68** |
